# Short- and long-term reconfiguration of rat prefrontal cortical networks following single doses of psilocybin

**DOI:** 10.1101/2024.12.10.627734

**Authors:** Ross J. Purple, Rahul Gupta, Christopher W. Thomas, Caroline T. Golden, Seán Froudist-Walsh, Matthew W. Jones

## Abstract

We quantify cellular- and circuit-resolution neural network dynamics following therapeutically relevant doses of the psychedelic psilocybin. Using chronically implanted Neuropixels probes, we recorded local field potentials (LFP) alongside action potentials from hundreds of neurons spanning infralimbic, prelimbic and cingulate subregions of the medial prefrontal cortex of freely-behaving adult rats. Psilocybin (0.3mg/kg or 1mg/kg i.p.) unmasked 100Hz high frequency oscillations that were most pronounced within the infralimbic cortex, persisted for approximately 1h post-injection and were accompanied by decreased net pyramidal cell firing rates and reduced signal complexity. These acute effects were more prominent during resting behaviour than during a sustained attention task. LFP 1-, 2- and 6-days post-psilocybin showed gradually-emerging increases in beta and low-gamma (20-60Hz) power, specific to the infralimbic cortex. These findings reveal features of psychedelic action not readily detectable in human brain imaging, implicating infralimbic network oscillations as potential biomarkers of psychedelic-induced network plasticity over multi-day timescales.

## INTRODUCTION

Increasing evidence suggests that psychedelic-induced ‘reconfiguration’ of brain connectivity patterns can culminate in lasting psychological changes, sparking substantial interest in psychedelic-based or -derived treatment of mental health problems (Carhart-Harris & Goodwin, 2017; Kočárová et al., 2021; Siegel et al., 2021). For example, in a recent phase IIb double blind trial, a single dose of synthesized crystalline polymorph COMP360 psilocybin was shown to significantly reduce depression scores over a period of 3 weeks in people with treatment-resistant depression (Goodwin et al., 2022). However, directly assessing cellular and circuit-level neural mechanisms remains challenging in human studies, particularly over multi-day timescales. Defining the molecular, cellular and circuit mechanisms driving psychedelic actions by translating between human brain imaging and animal neurophysiology therefore remains essential (Smausz et al., 2022).

Psychedelics like psilocybin directly modulate the serotonergic (5-HT) system. Psilocybin’s active metabolite, psilocin, is a partial agonist of 5-HT receptors (5HTRs), notably acting on 5-HT_2A_, 5-HT_1A_, and 5-HT_2C_ receptors (Erkizia-Santamaría et al., 2022; Tylš et al., 2023). 5-HT_2A_ and 5-HT_2C_Rs are G-protein coupled receptors (GPCRs) which, on activation, increase cell excitation through the phospholipase C signalling pathway. In contrast, 5-HT_1A_Rs are G_i_-GPCRs which inhibit the activity of adenylate cyclase, leading to decreased cell excitation (Andrade, 2011; Smausz et al., 2022). Both subtypes are widely expressed throughout the rodent, primate and human brain, including in the prefrontal cortex (Froudist-Walsh et al., 2023; Santana et al., 2004; Zilles & Palomero-Gallagher, 2017), which is reciprocally connected to serotonergic nuclei in the brainstem (Hajós et al., 1998). Immunohistochemical studies in rat have shown both 1A and 2A Rs are expressed in cortical pyramidal cells (1A: 60% of cells, 2A: ∼60-90% of cells in PFC) and GABAergic interneurons (1A: 25% of cells, 2A: 25% of cells in PFC) (Santana et al., 2004). Thus, while the psychedelic effects of drugs including psilocybin are generally thought to involve activation of 5-HT_2A_Rs (Halberstadt & Geyer, 2011), other 5HTR subtypes may contribute to therapeutic potential (Conn et al., 2024). Certainly, widespread 5HTR expression, bidirectional effects on cell excitation, and modulation of both glutamatergic and GABAergic cells makes pinpointing the actions of psilocybin a complex challenge.

### Acute effects of psychedelics on neural activity

Human fMRI and MEG analyses suggest that psychedelics increase neocortical neural signal diversity, an effect associated with desynchronisation within local network populations and desegregation of distributed brain networks (Carhart-Harris et al., 2016; Lord et al., 2019; Muthukumaraswamy et al., 2013; Schartner et al., 2017; Shinozuka et al., 2024). Invasive recordings of local field potentials (LFP) can be used to back-translate these analyses into animals, in which a concomitant decrease in distributed oscillatory power would be expected following systemic psychedelics. Indeed, psilocybin/psilocin has been shown to decrease low-frequency power in both humans (Kometer et al., 2015; Tyls et al., 2016) and rodents (Golden & Chadderton, 2022; Thomas et al., 2022; Vejmola et al., 2021). To date, very few studies have investigated the effects of psilocybin on the activity of individual neocortical neurons and their circuit interactions, though Golden & Chadderton (2022) showed that systemic injection of 2mg/kg psilocybin led to a net increase in population (mixed pyramidal and interneuron) firing rates in the cingulate cortex of head-fixed mice. Here, we used large-scale, high-density electrophysiology in freely-behaving rats to quantify how psilocybin affects the activities of different cells types at different doses across PFC subregions, and how synchrony between cells is affected.

### Lasting changes to network activity

Few studies have investigated the long-term effects of psilocybin on underlying brain activity. Skosnik et al. found preliminary evidence for a sustained increase in auditory stimulus train-evoked scalp EEG theta power 2 weeks after a single dose of psilocybin in people diagnosed with major depressive disorder. This increase in evoked theta was interpreted as a metric of synaptic plasticity and was also found to correlate with improved depression scores (Skosnik et al., 2023). Others have shown psilocybin can lead to lasting increases or decreases in human resting state functional connectivity (Barrett et al., 2020; McCulloch et al., 2022). On a cellular level, both rodent cell culture and in vivo studies have shown prolonged, psilocybin-induced increases in neuritogenesis and spinogenesis (Ly et al., 2018; Shao et al., 2021). Identifying the lasting effects of psilocybin on neural network dynamics could bridge between these human and animal findings, linking sustained restructuring of connectivity patterns with potential translational markers of therapeutic effects.

In this study, we therefore quantified the effects of individual systemic injections of therapeutically relevant doses of psilocybin on cellular and network activity recorded from the prefrontal cortex of freely behaving rats. We hypothesised that systemic psilocybin would have heterogenous effects on cell firing rates and alter coordinated activity between neural and interneuronal populations, modulating network state transitions, the complexity of the prefrontal dynamics and attenuating lower-frequency oscillations. In humans, psilocybin’s most pronounced acute and chronic effects on functional connectivity impact the default mode network (Siegel et al., 2024). We therefore hypothesised that psilocybin’s effects would be more prominent during resting than task-engaged behaviour and include lasting changes in network activity over the 1-week recording period.

## RESULTS

### Psilocybin did not significantly alter movement or behaviour

**Figure 1:**
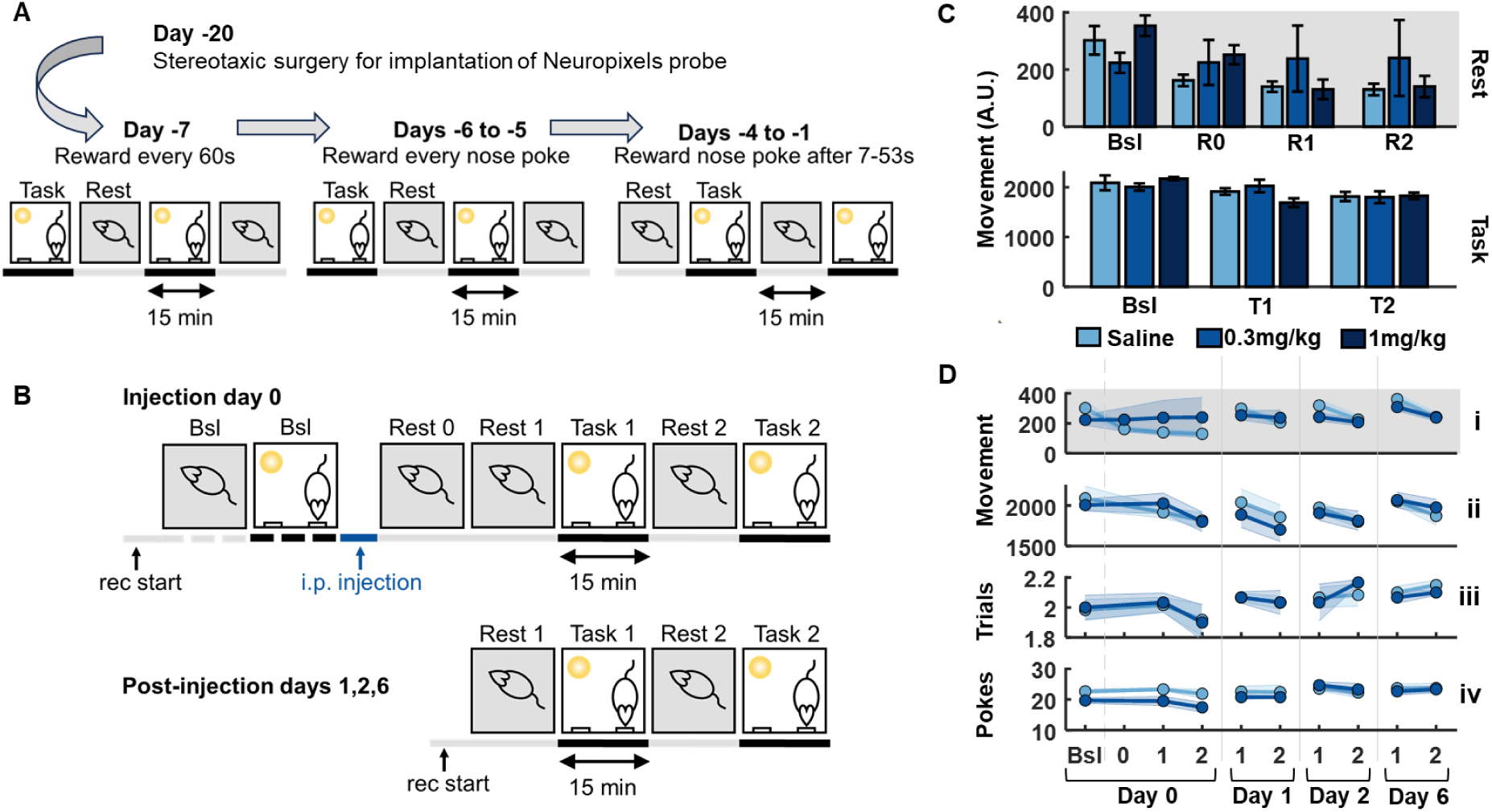
Activity and operant behaviour are not disrupted by systemic injection of 0.3mg/kg or 1mg/kg psilocybin. (A) Training on the operant task began approximately 2 weeks after chronic implantation of the Neuropixels probe into the medial prefrontal cortex (N=6 rats). Initial training sessions lasted for 1h: two 15min task blocks with the house light on and nose poke holes open, interleaved with two 15min rest blocks with the house light off and nose poke holes closed. On the first training day (day -7), ∼0.2ml of 30% sucrose solution liquid reward was dispensed automatically every 60s during the task block. On days -6/-5 a fixed ratio of one nose poke for one reward was applied. On days -4 to -1 animals performed the full operant task with the reward released upon nose poke after a random interval between 7-53s. The order of rest/task blocks was also switched. Rats were tethered from day -5 onwards to acclimatise to the electrophysiological recoding setup. **(B)** On the first injection day (Day 0) rats completed a 30min baseline session comprised of a 15min rest and 15min operant task block. Rats were then injected i.p. with either 0.3mg/kg psilocybin or saline vehicle (counterbalanced across animals), and again completed another operant task session with one initial 15min rest block (to account for initial drug effects) followed by a further four 15min blocks of alternating rest and task epochs (total 1hour 15min). Recordings were repeated on days 1, 2 and 6 with a 1h block of the operant task. 14 days after the first injection, rats received the alternative psilocybin/saline injection, and the protocol was repeated. A further 14 days after this, rats received a 1mg/kg injection of psilocybin, and the protocol was repeated. **(C)** Behavioural activity during day of injection. Bars show average video-recorded movement (a.u.) for each 15-minute block during rest (top) or task (bottom) blocks (Bsl=pre-injection baseline block). **(D)** Behavioural performance on the operant task across sessions during and following 0.3mg/kg psilocybin or saline. (i) Total activity (a.u.) during each rest block on the day of injection and days 1, 2 and 6 post-injection. (ii) Total activity (a.u.) during each task block across days. (iii) Average number of trials completed per minute for each task block. (iv) Average number of nose pokes to the rewarded hole per minute for each task block. Note that recordings were taken on the day of injection of 1mg/kg psilocybin injection, but not on days 1-6 after injection. N=4.

In this study we chronically implanted Neuropixels 1.0 probes and recorded neural activity across the medial prefrontal cortex (mPFC) of freely behaving rats. Rats performed an operant task in which nose-pokes were reinforced according to a random variable interval schedule in 15-minute blocks (task blocks) interleaved with 15-minute blocks of free behaviour (rest blocks) where no reinforcements were available, before and after administration of either 0.3mg/kg psilocybin, 1mg/kg psilocybin, or saline (Figure 1B). A comparison of video-recorded movement pre-to post-injection revealed similar activity levels after injection of either saline, 0.3mg/kg psilocybin, or 1mg/kg psilocybin during either task or rest blocks (ANOVA1: Rest blocks: F(5,18)=2.1, p=0.113; Task blocks: F(5,18)=2.53, p=0.067; Figure 1C), though the low frame-rate of video recordings meant head-twitch responses could not be reliably measured. On post-injection days 1, 2 and 6, performance of the operant task remained similar following 0.3mg/kg psilocybin and saline injections (Figure 1D), whether quantified using number of trials completed (RM-ANOVA: drug: F(1,6)=0.00, p=0.971, time: F(8,48)=0.46, p=0.880, drug x time: F(8,48)=0.20, p=0.990), number of nose pokes (RM-ANOVA: drug: F(1,6)=1.48, p=0.270, time: F(8,48)=3.15, p=0.006, drug x time: F(8,48)=1.69, p=0.125), or gross activity (RM-ANOVA: Rest blocks: drug: F(1,6)=0.00, p=0.957, time: F(9,54)=0.58, p=0.807, drug x time: F(9,54)=1.49, p=0.176; Task blocks: drug: F(1,6)=0.04, p=0.850, time: F(8,48)=1.40, p=0.222, drug x time: F(8,48)=0.91, p=0.518). Consistent performance on the task following vehicle and psilocybin injections therefore enabled analyses of drug effects during both task-focused and resting states.

### Psilocybin leads to acute emergence of a HFO within the infralimbic cortex

Local field potentials (LFP) within the infralimbic cortex revealed a sustained ∼100Hz high frequency oscillation (HFO) after 0.3mg/kg injection of psilocybin, evident during both task and rest periods (individual examples: Figure 2A). HFO power peaked during the second 15-minute rest block, lasting until at least 60 minutes post-injection (Figure 2B; RM-ANOVA: drug: F(1,6)=5.74, p=0.054, time: F(104,624)=5.26, p<0.001, drug x time: F(104,624)=2.63, p<0.001). Power spectra across 0-200Hz revealed a broadband decrease in power after 0.3mg/kg psilocybin, except at ∼100Hz. This effect was most discernible during rest blocks, compared to task blocks (Suppl. Fig. 2; ANOVA2: Rest blocks, time [pre vs post]: F(1,6408)=21.64, p<0.001, frequency: F(800,6408)=7.66, p<0.001, time x frequency: F(800,6408)=1.59, p<0.001; Task blocks, time [pre vs post]: F(1,6408)=19.24, p<0.001, frequency: F(800,6408)=15.67, p<0.001, time x frequency: F(800,6408)=0.93, p=0.925). Comparing spectral changes between psilocybin and saline revealed a significantly greater decrease post-injection of 0.3mg/kg psilocybin for frequencies between 20-80Hz (Figure 2C; ANOVA2: drug: F(2,8010)=67.04, p<0.001, frequency: F(800,8010)=7.94, p<0.001; drug x frequency: F(1600,8010)=0.77, p=1.000). Similar results were identified at 1mg/kg psilocybin albeit with a significant reduction in power between 3.5-4.5Hz and a shift in the peak of the HFO to a slower frequency of ∼93Hz (Figure 2C).

**Figure 2:**
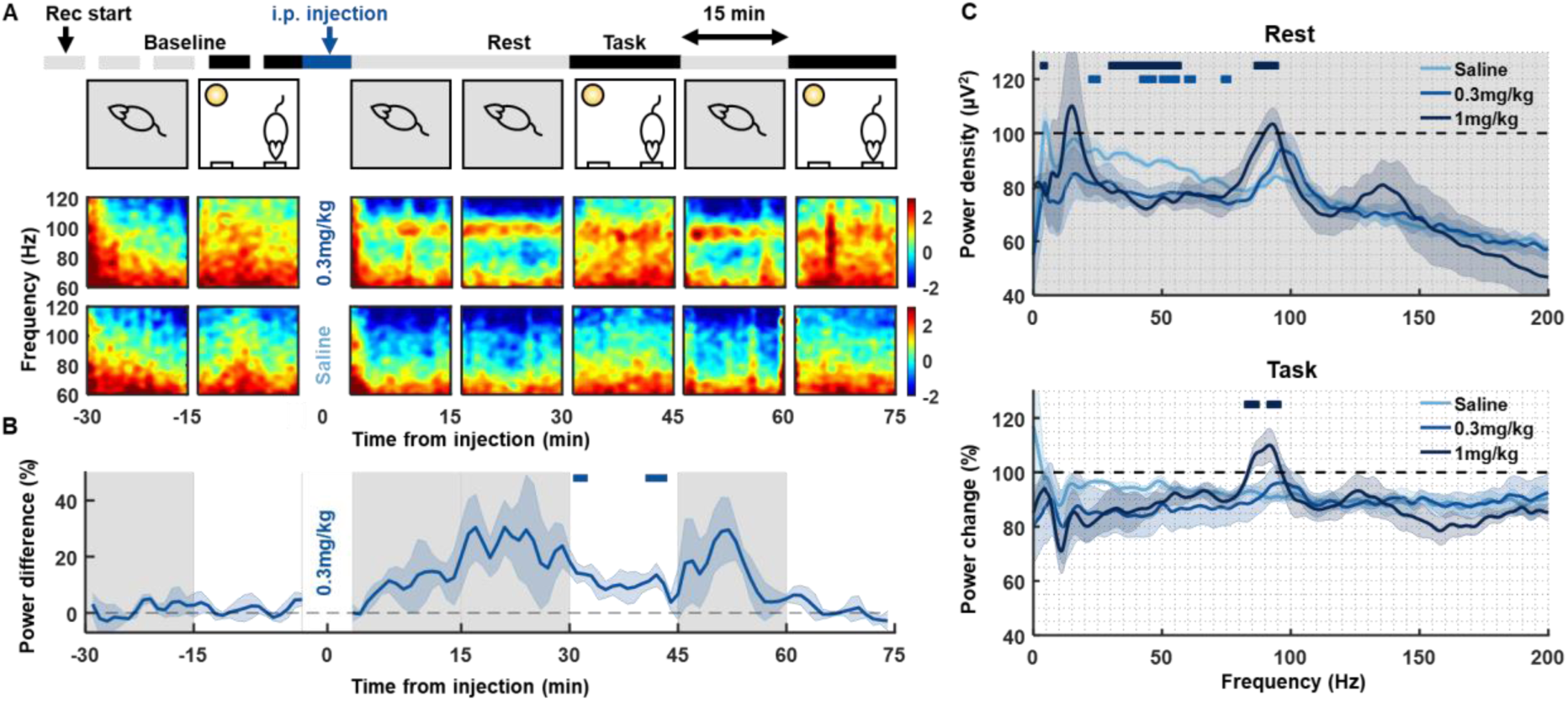
Psilocybin induces frequency-dependent changes in oscillations in medial prefrontal cortical local field potentials (LFP). **(A)** LFP spectrograms from a single recording channel in the infralimbic cortex of one animal aligned to rest and operant task 15min epochs pre- and post-injection of 0.3mg/kg psilocybin (top row spectrograms) and saline (bottom row). Note the sustained power around 100Hz during the rest block after injection of 0.3mg/kg psilocybin. **(B)** Average time course of ∼100Hz high frequency oscillation 1/f power difference at baseline and post-injection, shown as a difference in 1/f power difference between 0.3mg/kg psilocybin and saline. Grey shaded and white backgrounds delineate rest and operant task blocks respectively. Bars above trace mark times of significant difference to saline (partial-Bonferroni corrected post-hoc t-tests, p<0.05 **(C)** Average change in power density during the rest (top) and task (bottom) blocks from baseline (average from -30min to -15min pre-injection) to post injection (average from 15-30min and 45-60min post-injection) of saline (N=4), 0.3mg/kg psilocybin (N=5) and 1mg/kg psilocybin (N=4). Bars above power change spectra represent significant differences to saline for 0.3mg/kg psilocybin (lighter blue) and 1mg/kg (darker blue; partial-Bonferroni corrected post-hoc t-tests, p<0.05).

### Psilocybin-induced HFO center on the infralimbic cortex

To determine how the ∼100Hz HFO was anatomically distributed across the mPFC, spectral power was calculated during post-psilocybin injection rest blocks across all LFP channels (Figure 3A). For each of these outputs, a 1/f power difference was calculated by removing the signal between 80-110Hz and interpolating this gap to enable a calculation of the relative increase in ∼100Hz power (by taking the difference in power between the original signal and the interpolated signal at the peak frequency between 80-110Hz; Figure 3B). Mapping this result to the histological verification of the Neuropixels placement revealed a continuum of HFO power that was maximal within the infralimbic cortex (Figure 3C and 3D). The HFO 1/f power difference rapidly dissipated along the probe moving dorsally from the infralimbic cortex to the prelimbic and cingulate cortex, at which point it was no longer discernible. In one rat (rat , the Neuropixels probe was not placed in the infralimbic cortex and there was also no increase in the HFO 1/f power difference identified in this rat. While it was not possible to verify precisely which layers of the cortex each probe was placed into, the strongest increase in HFO power was clearly identified in the rats in which the probe was placed into the deeper layers of the cortex (rats 3, 4, 6 with the probe in layers III, V, VI). To quantify differences in HFO power across the probe, and across drug conditions, a generalised linear model was computed using a log-linked inverse gaussian distribution. In addition to main effects of distance along the probe (β=0.01, SE=0.01, t=2.53, p=0.011) and drug (β=0.20, SE=0.01, t=32.07, p<0.001), a significant interaction was also identified between distance and drug condition on HFO 1/f power difference (β=-0.05, SE=0.01, t=-20.95, p<0.001; Figure 3D).

**Figure 3:**
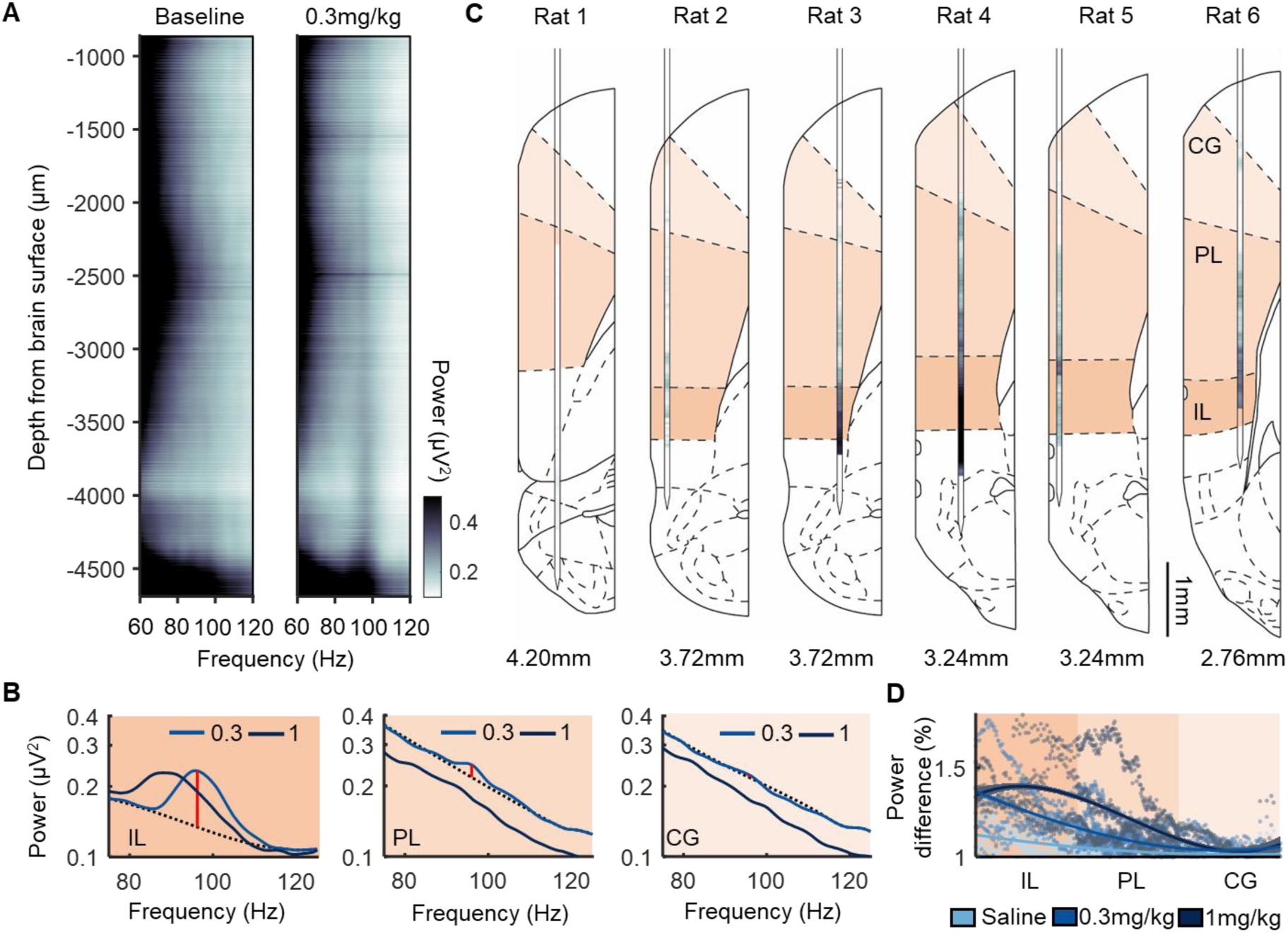
Sustained 80-110Hz oscillations (HFO) following psilocybin are most pronounced in infralimbic cortex. **(A)** LFP power between 60-120Hz across all electrode channels spanning the medial prefrontal cortex of one rat averaged across the 15-minute rest period pre-(left) and post-(right) injection of 0.3mg/kg psilocybin. **(B)** Power density from an LFP channel in the cingulate cortex (CG; top), prelimbic cortex (PL, middle) and infralimbic cortex (IL, bottom) during post-injection of 0.3mg/kg and 1mg/kg psilocybin. Dotted line denotes interpolated signal. Red line denotes HFO 1/f power difference between raw and interpolated signal. **(C)** HFO 1/f power difference across all recorded channels for each rat. Darker hues indicate greater difference in HFO power. Value below denotes anterior-posterior stereotaxic coordinate of the probe placement based on post-mortem reconstruction. **(D)** HFO 1/f power difference from all LFP electrodes in all rats as a function of their estimated dorsal-ventral location within the prefrontal cortex. Locations were normalised according to the size of each of the three regions assessed. Plot shows data from individual electrodes (dots) and polynomial fits (curves) for each drug condition with shaded areas depicting 95% prediction intervals.

### Psilocybin leads to an overall decrease in cell firing rates

Spike sorting analysis identified a median of 280 (range from 66-669) stable single units per animal during a single ∼2 hour recording session. There were no differences in the number of single units detected between each drug condition (Suppl. Fig. 3A). An assessment across time revealed that 0.3mg/kg of psilocybin led to a significant decrease in firing rates of pyramidal cells within the prelimbic cortex compared to saline. This effect was specific to the rest blocks and not apparent when the animal was actively engaged in the operant task (Figure 4A). No differences were identified between 0.3mg/kg psilocybin and saline within the cingulate or infralimbic cortex. In contrast, 1mg/kg psilocybin led to a significant decrease in firing rates in all three brain areas and was most apparent during the task blocks (Infralimbic: SRH: Drug: H(2,47460)=245.81, p<0.001, Time: H(104,47460)=553.96, p<0.001, Drug x Time: H(208,47460)=425.11, p<0.001; Wilcoxon: p<0.05 (at 1mg/kg only); Prelimbic: SRH: Drug: H(2,174510)=779.20, p<0.001, Time: H(104,174510)=954.46, p<0.001, Drug x Time: H(208,174510)=583.07, p<0.001; Wilcoxon: p<0.05 (at 0.3m/kg and 1mg/kg); Cingulate: SRH: Drug: H(2,22575)=452.03, p<0.001, Time: H(104,22575)=99.65, p=0.603, Drug x Time: H(208,22575)=309.08, p<0.001; Wilcoxon: p<0.05 (at 1mg/kg only)). No differences were identified in interneuron firing rates across time between conditions (Suppl. Fig. 3B).

A comparison of firing rate changes between rest and task blocks revealed that within the prelimbic cortex and infralimbic cortex, firing rates of pyramidal cells were significantly higher during rest blocks compared to task blocks during baseline conditions (Figure 4Bi; showing the difference between firing rates averaged during the baseline rest block OR baseline task block and firing rates averaged across baseline rest AND task blocks). No difference in firing rates was identified between task and rest blocks within the cingulate cortex, and there were no baseline differences in any brain region across drug conditions (SRH: Cingulate: Task: H(1,430)=0.33, p=0.568; Drug: H(2,430)=0, p=1.00; Task x Drug: H(2,430)=0.41, p=0.815; Prelimbic: Task: H(1,3324)=75.80, p<0.001; Drug: H(2,3324)=0, p=1.00; Task x Drug: H(2,3324)=13.04, p=0.002; Infralimbic: Task: H(1,904)=21.68, p<0.001; Drug: H(2,904)=0, p=1.00; Task x Drug: H(2,904)=2.94, p=0.230; Wilcoxon: p<0.05). The prelimbic difference between task and rest blocks remained significant post-injection of both saline and 1mg/kg psilocybin but was diminished after 0.3mg/kg psilocybin (Figure 4Bii; showing the difference between firing rates averaged during the post-injection rest blocks OR post-injection task blocks and firing rates averaged across baseline rest AND task blocks). All three brain regions showed a significant main effect of drug condition, post-injection (SRH: Cingulate: Task: H(1,430)=0.30, p=0.583; Drug: H(2,430)=23.74, p<0.001; Task x Drug: H(2,430)=0.79, p=0.674; Prelimbic: Task: H(1,3324)=9.99, p=0.002; Drug: H(2,3324)=34.68, p<0.001; Task x Drug: H(2,3324)=3.98, p=0.137; Infralimbic: Task: H(1,904)=3.71, p=0.054; Drug: H(2,904)=24.10, p<0.001; Task x Drug: H(2,904)=1.57, p=0.456; Wilcoxon: p<0.05).

A comparison of the scale of change in firing rates from baseline to post-injection revealed significantly greater change in firing rates during rest blocks, compared to task blocks after the 0.3mg/kg injection within the prelimbic cortex (Figure 4C; showing the difference between firing rates averaged during the post-injection rest blocks OR post-injection task blocks and firing rates averaged across baseline rest block OR baseline task block, respectively). This suggests the diminished separation in firing rates between rest and task blocks after 0.3mg/kg psilocybin is driven by the greater decrease in firing rates during rest blocks. Again, all three brain regions showed a significant main effect of drug condition (SRH: Cingulate: Task: H(1,430)=0. 30, p=0.87; Drug: H(2,430)=32.91, p<0.001; Task x Drug: H(2,430)=1.97, p=0.373; Prelimbic: Task: H(1,3324)=2.32, p=0.128; Drug: H(2,3324)=40.91, p<0.001; Task x Drug: H(2,3324)=18.54, p<0.001; Infralimbic: Task: H(1,904)=0.02, p=0.890; Drug: H(2,904)=27.56, p<0.001; Task x Drug: H(2,904)=1.04, p=0.594; Wilcoxon: p<0.05). Finally, an overall assessment of differences between brain regions in the pre-to post-drug change revealed no differences during rest blocks (SRH: Rest blocks: Drug: H(2,2329)=28.13, p<0.001, Region: H(2,2329)=5.44, p=0.066, Drug x Region: H(4,2329)=4.97, p=0.291). However, this effect was significant during task blocks with the infralimbic cortex showing the greatest decrease in firing rates compared to both prelimbic and cingulate cortices (SRH: Task blocks: Drug: H(2,2329)=44.34, p<0.001, Region: H(2,2329)=9.31, p=0.010, Drug x Region: H(4,2329)=9.27, p=0.055; Wilcoxon: cingulate vs prelimbic, z=1.47, p=0.140; cingulate vs infralimbic, z=2.71, p=0.007, prelimbic vs infralimbic, z=2.14, p=0.033).

**Figure 4:**
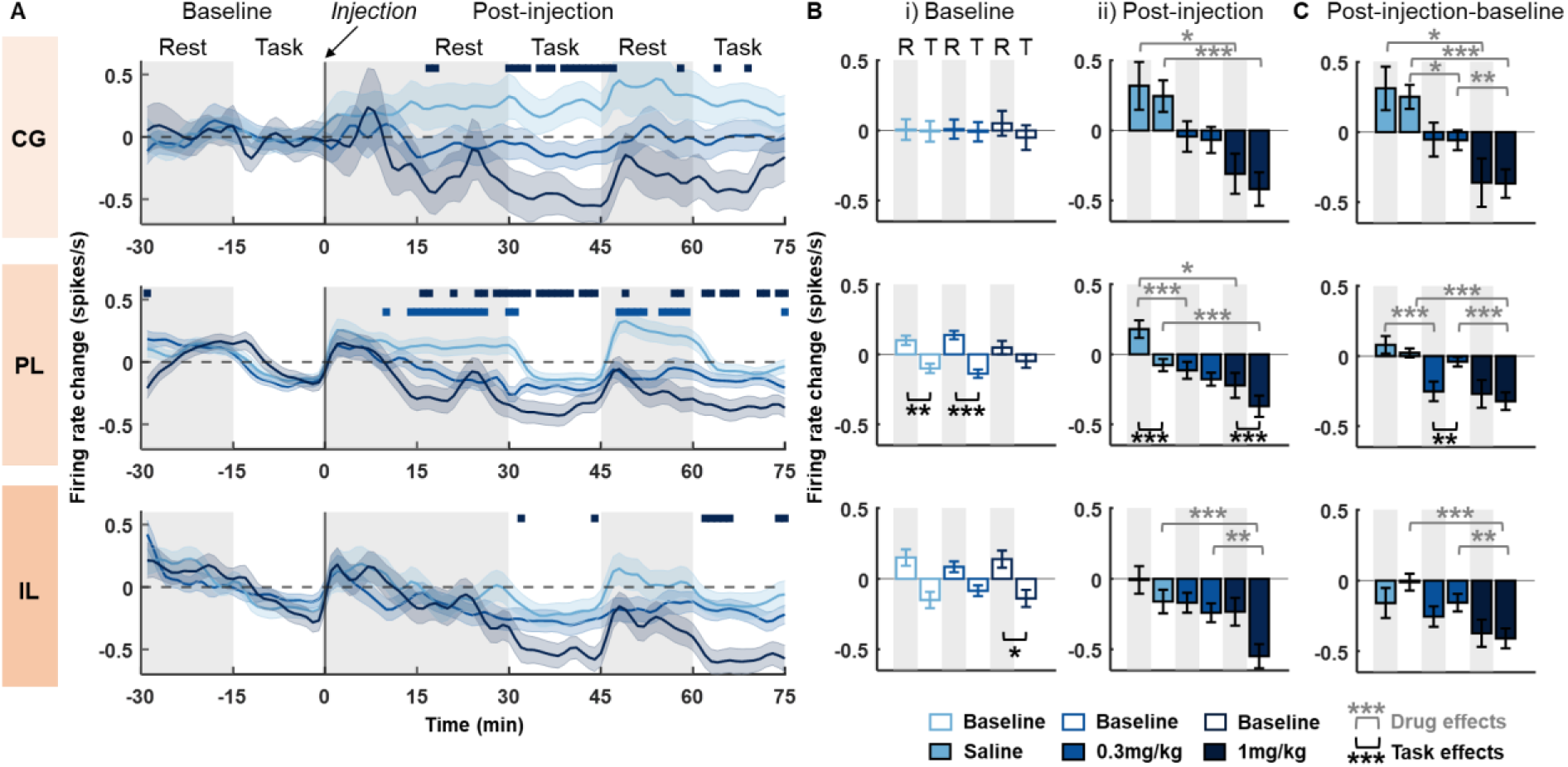
Psilocybin leads to a dose dependent net decrease in pyramidal cell firing rates that differs across brain regions and behavioural state. **(A)** Time-course of change in firing rates of pyramidal cells in the cingulate cortex (CG; top), prelimbic cortex (PL; middle) and infralimbic cortex (IL; bottom) relative to pre-injection baseline (injection at time 0). Firing rates are shown as a difference from the average firing rates during the 30-minute pre-injection baseline. Bars above the rate change indicate significant differences of 0.3mg/kg (light blue) and 1mg/kg (dark blue) versus saline vehicle injection (post-hoc Bonferroni corrected Wilcoxon rank sum test: p<0.05). Grey shaded and white backgrounds delineate rest and operant task blocks respectively. **(B)** Average change in firing rates during rest (R) and task (T) blocks from pyramidal cells within the CG (top row), PL (middle row) and IL (bottom row). (i) average firing rate change during individual 15-minute rest and task blocks at baseline (shown as a difference from average firing rate across the whole 30-minute baseline period). (ii) average firing rate change from baseline to post-injection. **(C)** Difference in firing rate change between post-injection blocks (shown in Bii) and baseline blocks (shown in Bi), calculated separately for rest (post injection rest – baseline rest) and task (post-injection task – baseline task). Asterisks denote significant post-hoc Bonferroni corrected Wilcoxon rank sum or signed rank differences between rest and task blocks (signed rank; black) or between drugs (rank sum; grey): ***=p<0.001, **=p<0.01, *=p<0.05. Shaded bands in (A) and error bars in (B/C) denote SEM.

Overall, firing rates for both pyramidal cells and interneurons showed diverse changes across the mPFC for all injection conditions (Figure 5A). To further explore regional differences, we next compared the proportion of pyramidal cells and interneurons across regions that showed significant changes in firing rates pre-to post-injection. This analysis revealed a significantly higher proportion of pyramidal cells within the infralimbic and prelimbic cortex that had either decreased or no change in firing rates post-injection of 0.3mg/kg psilocybin, compared to saline (Figure 5B; chisq: saline vs 0.3mg vs 1mg; infralimbic decrease: X^2^=11.98, p=0.003; infralimbic no change: X^2^=10.13, p=0.006; infralimbic increase: X^2^=10.03, p=0.007; prelimbic decrease: X^2^=26.05, p<0.001; prelimbic no change: X^2^=18.61, p<0.001; prelimbic increase: X^2^=8.60, p=0.014; post-hoc chisq, p<0.05). There were no differences in the proportion of cells changing firing rates in the cingulate cortex between 0.3mg/kg psilocybin and saline (chisq: saline vs 0.3mg vs 1mg; cingulate decrease: X^2^=11.88, p=0.003; cingulate no change: X^2^=2.17, p=0.338; cingulate increase: X^2^=7.94, p=0.019; post-hoc chisq, p>0.05). Following 1mg/kg psilocybin, the proportion of cells with decreased post-injection firing rates was significantly higher in all three brain regions, in comparison to saline (post-hoc chisq, p<0.05). In general, interneurons also showed a greater proportion of cells with decreased firing rates in both infralimbic and prelimbic cortices, but this only reached significance in the infralimbic cortex at 0.3mg/kg psilocybin and in the prelimbic cortex at 1mg/kg psilocybin in comparison to saline, after controlling for multiple testing (chisq saline vs 0.3mg vs 1mg; infralimbic decrease: X^2^=7.15, p=0.028; infralimbic no change: X^2^=0.43, p=0.808; infralimbic increase: X^2^=5.08, p=0.079; prelimbic decrease: X^2^=8.72, p=0.013; prelimbic no change: X^2^=4.47, p=0.107; prelimbic increase: X^2^=1.03, p=0.598; cingulate decrease: X^2^=1.57, p=0.456; cingulate no change: X^2^=6.12, p=0.047; cingulate increase: X^2^=2.91, p=0.233; post-hoc chisq, p<0.05).

**Figure 5:**
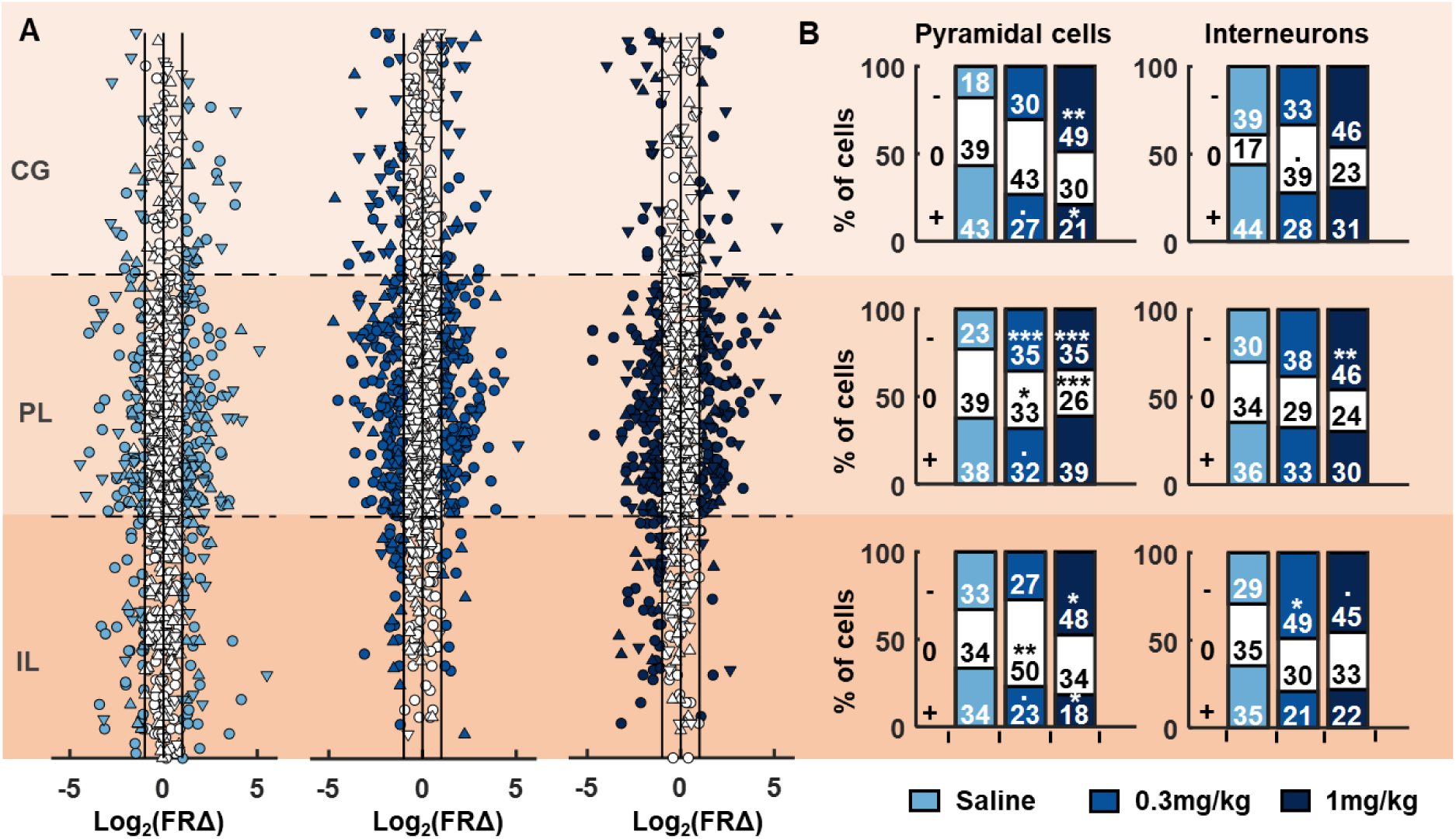
Pyramidal cells and interneurons of the medial prefrontal cortex respond to systemic psilocybin with heterogeneous changes in firing rate. **(A)** Individual cell firing rate changes during pre-to post-injection rest blocks sorted according to estimated dorsal-ventral location within the prefrontal cortex. Each of the three brain regions (IL, PL, CG) were normalised in size with approximate location of each cell determined by proximity to electrodes along the probe. Cells coloured in white showed either less than a twofold change in firing rates, or the change did not reach significance. Distributions are shown separately for saline (left), 0.3mg/kg psilocybin (middle) and 1mg/kg psilocybin (right). Circles denote putative pyramidal cells, triangles pointing up denote narrow-waveform interneurons, triangles pointing down denote wide-waveform interneurons. **(B)** Proportion of pyramidal cells (left) and narrow-waveform interneurons (right) showing a significant decrease (-), increase (+) or no change (0) from baseline to post-injection of psilocybin or saline within each brain region during rest blocks. Asterisks denote significant post-hoc chi-squared test differences in the proportion of cells between saline and 0.3mg/kg or 1mg/kg psilocybin; ***=p<0.001, **=p<0.01, *=p<0.05, .=p<0.05 but did not survive Bonferroni multiple comparison correction.

### Psilocybin enhances cell pair cross-correlations

We next assessed the degree to which synchronous cell firing was modulated by psilocybin. An analysis of cell pair cross-correlations revealed a marked increase in co-activity between pairs of pyramidal cells and pyramidal-interneuron pairs within the infralimbic cortex, after injection of both 0.3mg/kg and 1mg/kg psilocybin (Figure 6A). This increase from baseline to post-injection was significantly greater in comparison to the effects of the saline injection (Figure 6B, left; KW: Pyr-Pyr: X^2^(2,3086)=18.78, p<0.001; Pyr-Int: X^2^(2,2412)=27.45, p<0.001; Wilcoxon: p<0.05). No changes were identified for pairs of interneurons (KW: Int-Int: X^2^(2,459)=0.82, p=0.663). For cell pairs within the prelimbic cortex, a significant increase in cross-correlated activity between pairs of pyramidal cells was identified at 1mg/kg but not 0.3mg/kg psilocybin, in comparison to saline (Figure 6B, right & Suppl.Fig. 4; KW: Pyr-Pyr: X^2^(2,22239)=56.04, p<0.001; Wilcoxon: p<0.05). No differences were identified for pyramidal-interneuron pairs (KW: Pyr-Int: X^2^(2,11648)=3.98, p=0.137). In contrast, a significant decrease was found for pairs of interneurons at 1mg/kg psilocybin in comparison to saline (KW: Int-Int: X^2^(2,1475)=21.18, p<0.001; Wilcoxon: p<0.05). No differences were found for any combination of cell-type pairs between the infralimbic and prelimbic cortices (Suppl. Fig. 5; KW: Pyr-Pyr: X^2^(2,11712)=1.58, p=0.454; Int-Int: X^2^(2,1174)=2.81, p=0.245; Pyr-Int: X^2^(2,7600)=0.60, p=0.742; N.B. not enough cells were identified from the cingulate cortex to carry out analysis in this region). Cross-correlated activity during pre-injection baseline was not significantly different between 0.3mg/kg psilocybin, 1mg/kg psilocybin and saline (infralimbic, KW: Pyr-Pyr: X^2^(2,3086)=2.23, p=0.327; Int-Int: X^2^(2,459)=0.61, p=0.739; Pyr-Int: X^2^(2,2410)=5.62, p=0.060).

**Figure 6:**
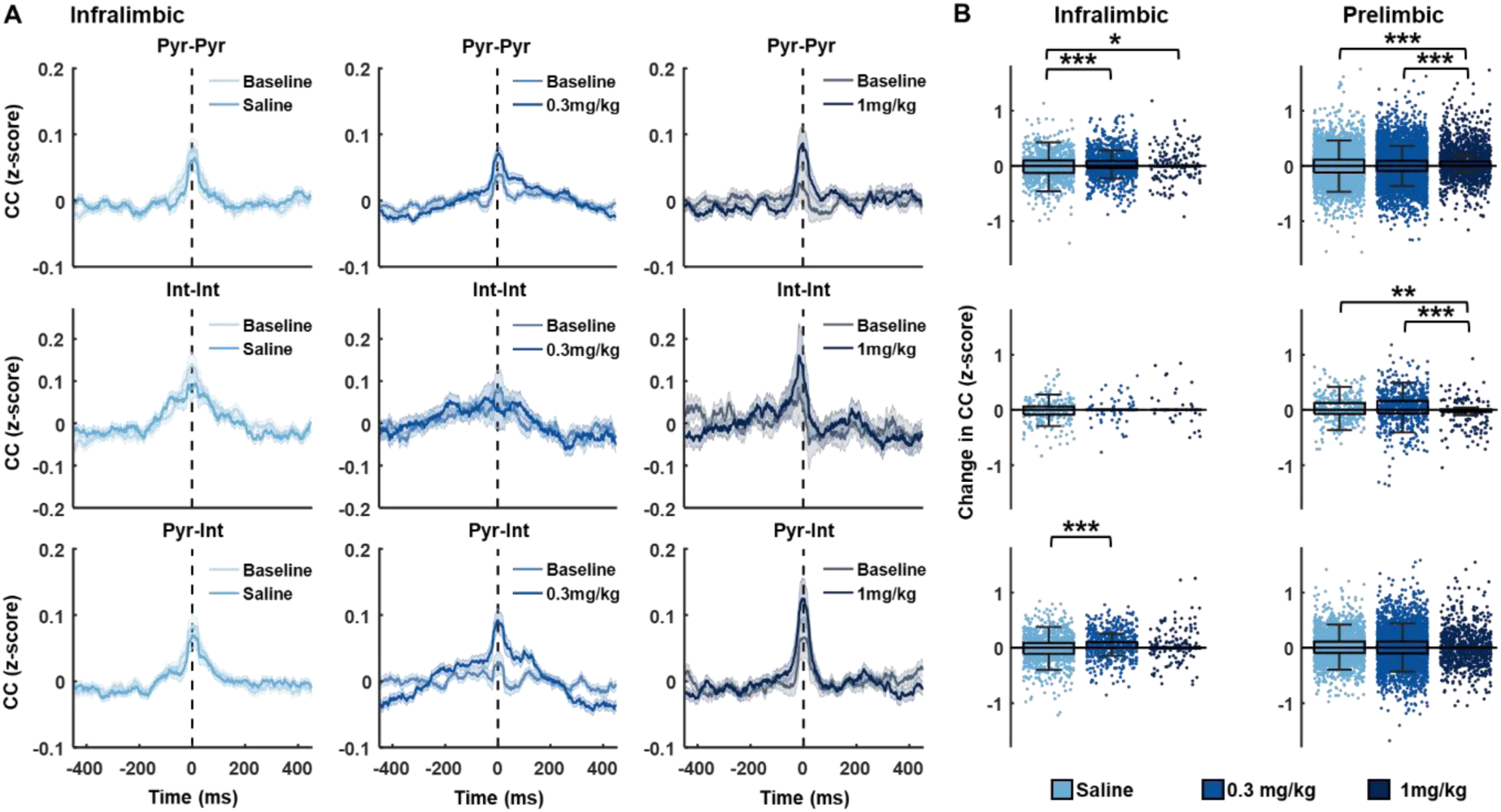
Psilocybin augments cross-correlated activity between pyramidal cells and interneurons in infralimbic cortex. **(A)** Cross-correlated activity (2ms bin size) between cell pairs within the infralimbic cortex at baseline rest, and post-injection of saline (left, grey), 0.3mg/kg psilocybin (middle, light blue) and 1mg/kg psilocybin (right, dark blue). Shown are pyramidal-pyramidal (pyr-pyr), interneuron-interneuron (int-int) and pyramidal-interneuron (pyr-int) cell pair correlations. **(B)** Change in cross-correlated activity from baseline to post-injection rest blocks for pyr-pyr (top), int-int (middle) and pyr-int (bottom) cell pairs within the infralimbic cortex (left) or prelimbic cortex (right). A significant increase in cross-correlated activity was identified for pyr-pyr and pyr-int cell pairs within the infralimbic cortex after injection of 0.3mg/kg psilocybin, compared to saline (Bonferroni corrected post-hoc t-tests, p<0.05). Within the prelimbic cortex a significant increase in cross-correlated activity was identified for pyr-pyr and a significant decrease was identified for int-int cell pairs after injection of 1mg/kg psilocybin (Bonferroni corrected post-hoc t-tests, p<0.05).

### Psilocybin causes less complex mPFC activity with a deeper energy landscape

Since the single units recorded in the mPFC constitute a subset of larger local neuronal populations, we next considered their activities collectively as a network, representative of mPFC dynamics. The time-binned spikes from all units were binarized and the mean-squared displacement (MSD) was calculated to describe the scale of network state transition across time (Figure 7A). This analysis revealed a significant increase in MSD distributions from pre-to post-injection of saline, in contrast to a significant decrease after both 0.3mg/kg and 1mg/kg psilocybin (Figure 7B; KW: Rest blocks: saline: X^2^(1,359598)=3357.1 p<0.001, 0.3mg/kg: X^2^(1,359598)=46604.58 p<0.001, 1mg/kg: X^2^(1,359598)=26416.76 p<0.001). Comparing this change across conditions from pre-to post-injection, confirmed a significant decrease in MSD distributions for both 0.3mg/kg and 1mg/kg psilocybin compared to saline, and a further decrease between 0.3mg/kg and 1mg/kg psilocybin (Figure 7C; KW: Rest blocks: saline vs 0.3mg/kg: X^2^(1,359598)=77897.8 p<0.001; saline vs 1mg/kg: X^2^(1,359598)=47149.3 p<0.001, 0.3mg/kg vs 1mg/kg: X^2^(1,359598)=765.34, p<0.001, Task blocks: saline vs 0.3mg/kg: X^2^(1,359598)=6479.48 p<0.001; saline vs 1mg/kg: X^2^(1,359598)=28210.89 p<0.001, 0.3mg/kg vs 1mg/kg: X^2^(1,359598)=10953.73 p<0.001). While the effect of the drug on decreasing MSD was significant for both rest and task blocks, the drug effect was of large effect size during rest periods (Cohen’s d, Rest blocks: saline vs 0.3mg/kg: d=1.04, saline vs 1mg/kg: d=0.81, 0.3mg/kg vs 1mg/kg: d=0.06), and small effect size during task periods (Task blocks: saline vs 0.3mg/kg: d=0.27, saline vs 1mg/kg: d=0.54, 0.3mg/kg vs 1mg/kg: d=0.31). Additionally, for each drug, the post-drug MSD distributions for rest and task blocks were significantly different (Rest vs Task: sal X^2^(1,359598)=3532.98 p<0.001, 0.3mg/kg X^2^(1,359598)=33825.29 p<0.001, 1mg/kg X^2^(1,359598)=1287.25 p<0.001).

The network state transitions shaped by the energy landscape also underlies the information content of the network dynamics. Frequent transitions across a variety of states can lead to higher information richness and entropy of the network activity (Singleton et al., 2022). Lempel-Ziv complexity of the binarized neural spiking activity can be used as a measure of entropy (Szczepański et al., 2004). Therefore, we computed the LZ complexity of binned spiking activity of the individual units. While there were no significant differences pre-to post-drug injections under saline, a noticeable decrease in complexity was observed after both 0.3mg/kg and 1mg/kg psilocybin (Figure 7D; PT, one-tailed: saline: p=0.3131, 0.3mg/kg: p=0.0280, 1mg/kg: p=0.0148; PT, two-tailed: saline: p=0.6260, 0.3mg/kg: p=0.0595, 1mg/kg: p=0.0304). To measure the post-drug change in complexity during rest and task blocks, we pair-wise subtracted the units’ LZ complexity values during the pre-drug rest/task blocks from that during the post-drug rest/task blocks. 0.3mg/kg psilocybin led to a significant decrease in complexity compared to saline during rest, but not task blocks (Independent 2-sample t-test: rest p<0.001, task p=0.3096). 1mg/kg psilocybin led to a significant decrease in complexity compared to saline during both rest and task blocks (Figure 7E; Independent 2-sample t-test: rest p<0.001, task p<0.001). During the task block, 1mg/kg led to a significant complexity decrease compared to 0.3mg/kg, but not during rest blocks (Independent 2-sample t-test: rest p=0.5198, task p<0.001). We observed a significant effect of task (vs rest) on LZ complexity for 0.3 mg/kg psilocybin, but not for saline or 1mg/kg psilocybin (paired t-test: saline p=0.063, 0.3mg/kg: p<0.001, 1mg/kg: p=0.692). Collectively, these results suggest that psilocybin leads to ordered and less chaotic mPFC activity, by deepening the underlying energy landscape of mPFC dynamics, restricting wide and frequent network state transitions.

**Figure 7:**
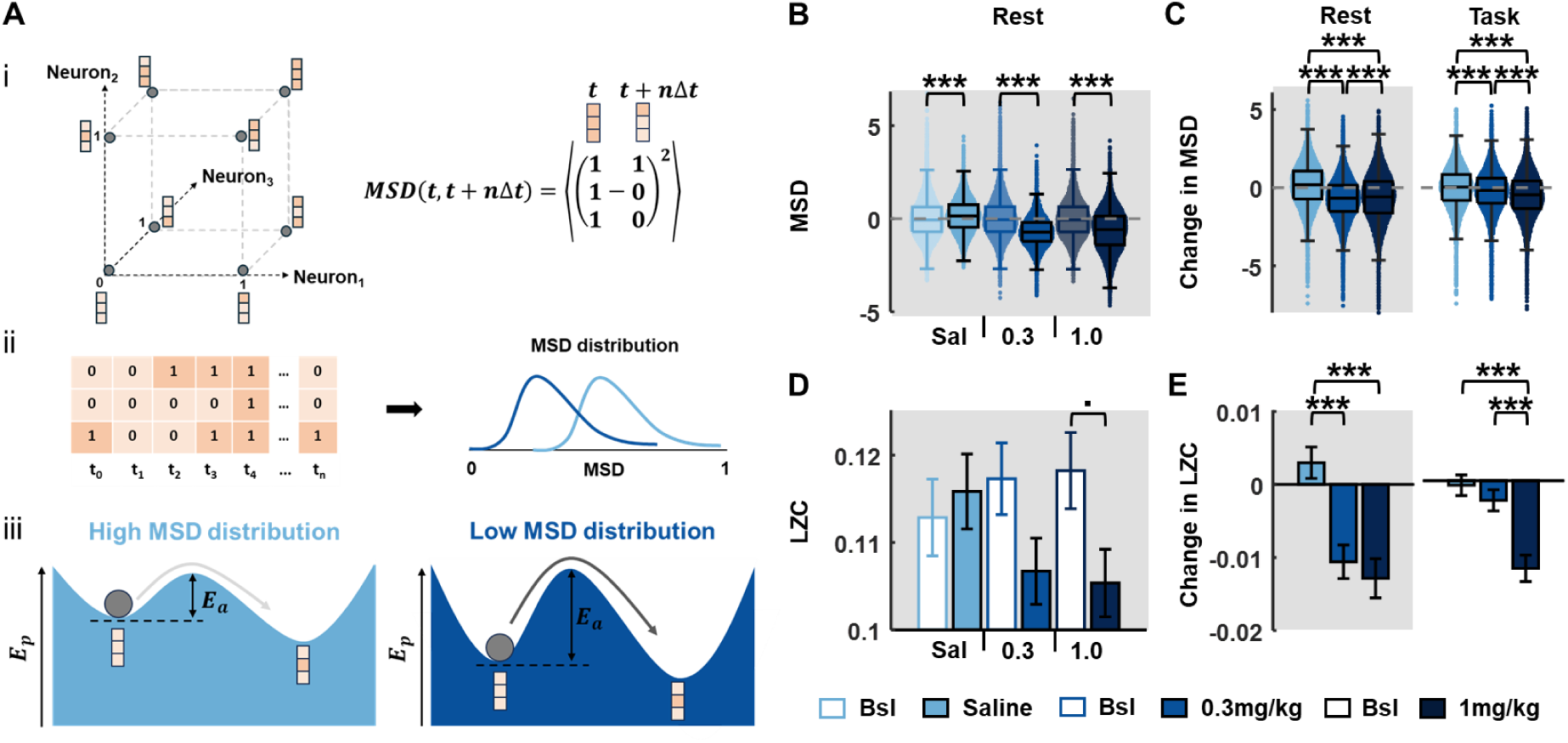
Psilocybin leads to decreased mean square displacement (MSD) of neural spiking activity and decreased Lempel-Ziv (LZ) complexity. **(A)** Distribution of mean-squared displacements (MSDs) of binarized network neural spiking states explains the energy landscape of network dynamics: The spiking activities of recorded single units in the mPFC can be used to describe the binary state transitions of a representative network of the units: (i, left) An example three-neuron network is shown with possible states visualised as the vertices of a unit cube. The possible 3-bit binary states, where 0 (light-orange) represents no spikes in the time bin, and 1 (dark-orange) represents spiking. The axes represent the binary values of the individual neurons. The system of neural network (dark-grey ball) transitions from one state to another with time. (i, right) The MSD is a Euclidean jump distance between any two network states over a discrete time-step Δ*t*. It is the squared difference between the network states at time *t* and *t* + Δ*t*, averaged 〈. 〉 over the number of neurons. (ii) The temporal dynamics of network is shown as consecutive 3-bit states at discrete time points, t1 to tn, derived from the time-binning (Δ*t*) of the units’ spike activities. At every time point, an average MSD of the network transitions over next 1 second is computed. This provides a distribution of MSDs as the statistical characteristics of network transitions across the entire time duration. MSD distributions centred on lower values reflect restricted state transitions, while higher values indicate relatively free network state transitions. (iii) Schematic descriptions of the potential energy landscapes, Ep, for the low and high MSD distributions are shown. Shallower landscapes (left) require less activation energy leading to more frequent state transitions (light grey arrow) and a high MSD distribution. The network activity (dark-grey ball) that is sitting in a deep basin of attraction (right) requires large activation energy, Ea, to jump to neighbouring states, causing less-frequent transitions (dark grey arrow) and low MSD distribution. **(B)** MSD from all single units across pre-and post-injection rest blocks for each drug condition, indicating a decrease in MSD (more restricted transitions) under psilocybin. **(C)** Change in MSD from pre-to post-injection rest blocks (left) and task blocks (right), across drug conditions, indicating a dose-dependent decrease in MSD with psilocybin. **(D)** LZ complexity from all single units across pre- and post-injection rest blocks for each drug condition. **(E)** Change in LZ complexity from pre-to post-injection rest blocks (left) and task blocks (right), across drug conditions, indicating a decrease in LZ complexity with psilocybin (0.3 and 1 mg/kg) in both rest and task contexts. Error bars denote SEM. Asterisks denote significant post-hoc t-test Bonferroni corrected differences between saline and 0.3mg/kg or 1mg/kg psilocybin; ***=p<0.001, **=p<0.01, *=p<0.05, .=p<0.05 but did not survive Bonferroni multiple comparison correction.

### Long-term effects of psilocybin on coordinated oscillations in the infralimbic cortex

To identify potential long-term effects of low-dose psilocybin, recordings on Day 0 were compared with drug-free activity on post-injection days 1, 2 and 6 post-injection. On Day 0, 0.3mg/kg psilocybin induced a significant decrease in 20-80Hz power during rest (but not task) blocks within the infralimbic cortex (Figure 8A; ANOVA2: Rest blocks, Day 0: Frequency: F(800,4806)=17.24, p<0.001; Drug: F(1,4806)=13.46, p<0.001; Frequency x Drug: F(800,4806)=62.43, p=0.003; Task blocks, Day 0: Frequency: F(800,4806)=1.77, p<0.001; Drug: F(1,4806)=0.05, p=0.832; Frequency x Drug: F(800,4806)=0.58, p=1.000). Both rest and task blocks also showed an increase at ∼100Hz, though this was only statistically different to saline during task blocks (Figure 8A, see Suppl. Fig.6 for separated results from saline and psilocybin). This Infralimbic HFO power difference was an acute effect of psilocybin and absent from day 1 onwards. However, across days there was a gradual increase in power between 20-60Hz during both rest and task blocks (Figure 8A; ANOVA2: Rest Day 6: Frequency: F(800,4806)=7.4, p<0.001; Drug: F(1,4806)=177, p<0.001; Frequency x Drug: F(800,4806)=1.24, p<0.001; Task Day 6: Frequency: F(800,4806)=0.48, p=1.000; Drug: F(1,4806)=28.94, p<0.001; Frequency x Drug: F(800,4806)=1.31, p<0.001). This psilocybin-induced evolution of 20-60Hz power across days was not evident for LFPs in the prelimbic or cingulate cortices (Suppl. Fig.7).

On Day 0, a broad increase in coherence between pairs of LFP channels was identified across all frequencies following 0.3mg/kg psilocybin during rest blocks but a significant decrease in coherence >100Hz during task blocks (Figure 8B, shown for IL-PL, with similar findings for IL-CG and PL-CG; ANOVA2: IL-PL, Rest Day 0: Frequency: F(800,4806)=7.86, p<0.001; Drug: F(1,4806)=2176.07, p<0.001; Frequency x Drug: F(800,4806)=0.73, p=1.000; IL-PL, Task Day 0: Frequency: F(800,4806)=1.05, p=0.186; Drug: F(1,4806)=427.64, p<0.001; Frequency x Drug: F(800,4806)=0.28, p=1.000). Similar results were found for both power and coherence at 1mg/kg dose of psilocybin in comparison to saline (Suppl. Fig.8). These coherence patterns evolved across Days 1-6, gradually decreasing at frequencies below 100Hz across days and culminating in a broadband decrease in coherence by day 6 (Figure 8B, shown for IL-PL LFPs; ANOVA2: Rest Day 6: Frequency: F(800,4806)=2.38, p<0.001; Drug: F(1,4806)=1744.14, p<0.001; Frequency x Drug: F(800,4806)=0.34, p=1.000; Task Day 6: Frequency: F(800,4806)=0.61, p=1.000; Drug: F(1,4806)=1809.94, p<0.001; Frequency x Drug: F(800,4806)=0.59, p=1.000; ANOVA2 results for day 1 and 2 can be found in Suppl. Table 1).

**Figure 8:**
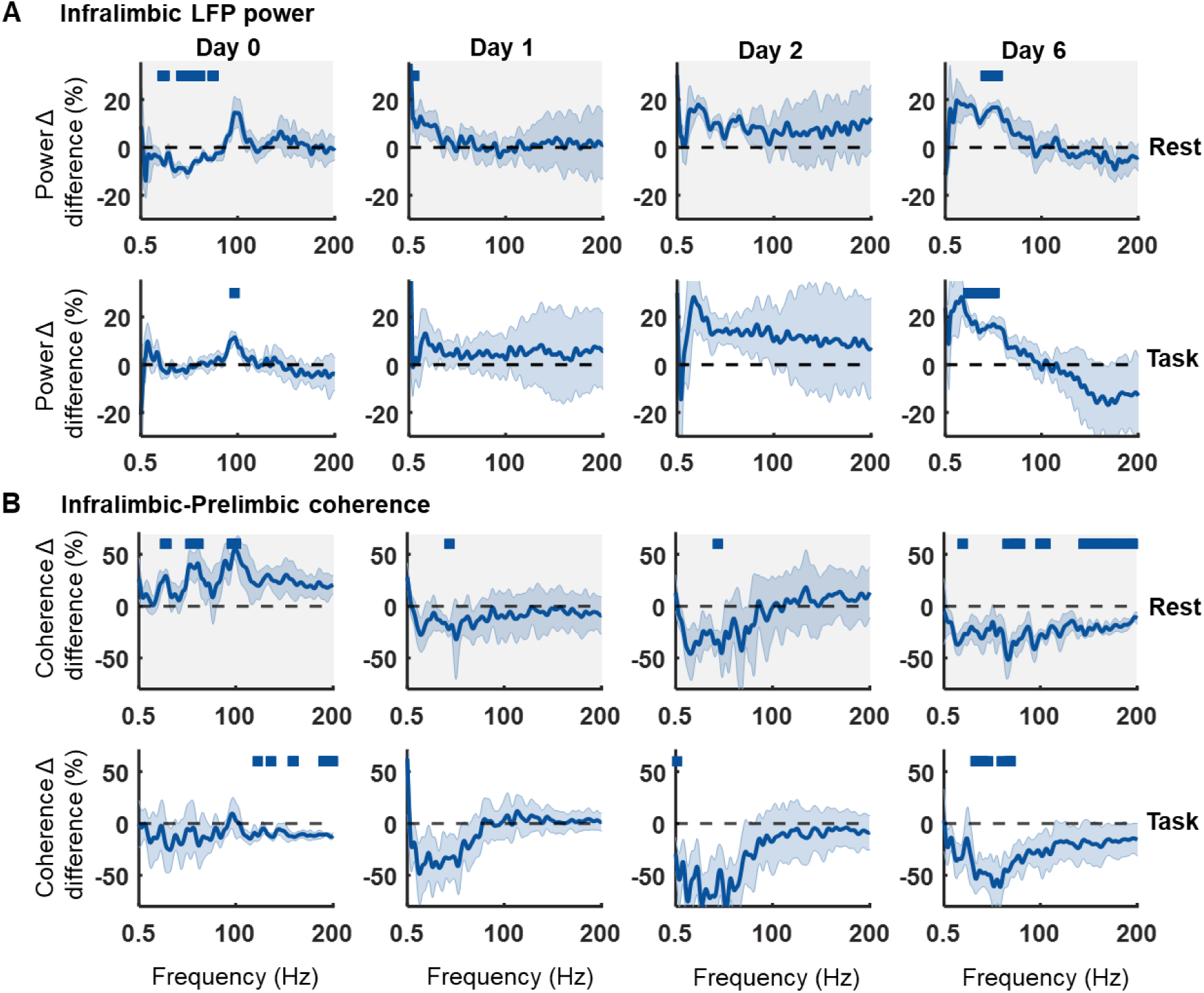
A single injection of psilocybin induces long-term changes in LFP power in infralimbic cortex across 6 days post-injection. **(A)** Difference between 0.3mg/kg psilocybin in comparison to saline in the change in power density from baseline to post injection during rest (top, grey) and task (bottom, white) blocks, on the day of injection (Day 0) and subsequent days 1, 2 and 6 (all compared to day 0 baseline). **(B)** Difference between 0.3mg/kg psilocybin in comparison to saline in the change in infralimbic-prelimbic coherence from baseline to post injection across days during rest (top, grey) and task (bottom, white) blocks (all compared to day 0 baseline). Bars indicate significant differences between 0.3mg/kg psilocybin and saline (partial-Bonferroni corrected post-hoc t-tests, p<0.05). Shaded bands denote SEM.

## DISCUSSION

We show that systemic injections of clinically relevant doses of psilocybin, reconfigure network activity across the rat medial prefrontal cortex, with most effects centred on the infralimbic subregion during resting behaviour. Psilocybin induced a mix of firing rate changes across different cells but led to dose-dependent net decreases in the firing rates of both pyramidal cell and interneuron populations, particularly during rest periods when animals were disengaged from the task. This net suppression of spiking activity was accompanied by significant increases in cross-correlated activity between pyramidal cell and pyramidal-interneuron pairs specifically within the infralimbic cortex, and by a slowing of neural dynamics and a reduction in its complexity. At the network level, local field potentials revealed the infralimbic cortex as the epicentre of an acute ∼100Hz high-frequency oscillation, and the subsequent emergence over several days of increased 20-60Hz power and decreased intra-cortical coherence.

### Acute effects of psilocybin on oscillatory network activity

Psilocybin has previously been shown to acutely decrease low-frequency network oscillations, potentially reflecting desynchronisation of cortical networks, decreased top-down inhibitory control and consequent modulation of distributed cognitive processing (Carhart-Harris & Goodwin, 2017; Smausz et al., 2022). While 2mg/kg psilocybin decreased delta power within the cingulate cortex in mice (Golden & Chadderton, 2022), we show only marginal decreases in low frequency power immediately following the 1mg/kg dose. We also identified acute decreases in LFP gamma power after both 0.3mg/kg and 1mg/kg psilocybin. Our results are consistent with psilocybin-induced broadband decreases in power over frontal areas previously shown in rats (Vejmola et al., 2021) and humans (Barnett et al., 2020; Muthukumaraswamy et al., 2013).

How these mPFC low-frequency and gamma changes relate to interactions across longer-range brain networks remains to be established. Analyses of psilocybin’s effects on distributed functional connectivity based on extracranial recordings of human MEG (Barnett et al., 2020) and rat EEG (Vejmola et al., 2021) have tended to identify decreased functional connectivity at low frequencies. In contrast, we identified acute, broadband increases in LFP coherence between subregions of the mPFC after 0.3mg/kg psilocybin. Different species, doses and methodologies make direct comparison with other studies challenging. However, binding occupancy of psilocybin to 5-HT2ARs is around 21% at 0.2mg/kg and 41% at 1mg/kg in rats (Kiilerich et al., 2023), hence the increased broadband intracortical coherence following the lower 0.3mg/kg dose in our study could reflect modulation of more frontally-localized neural circuits at lower receptor occupancies.

Consistent with this suggestion, psilocybin’s acute induction of sustained, 100Hz HFO did appear to be localized to the infralimbic cortex. These HFO are reminiscent of those induced by dissociative NMDA receptor channel blockers including ketamine, MK801 and phencyclidine (Hunt et al., 2019; Phillips et al., 2012). Infralimbic HFO may arise from elsewhere; afferent projections with greatest specificity to the IL, in comparison to PL and CG, include the amygdalo-piriform transition zone and the horizontal limb of the diagonal band of Broca (DBh) in the basal forebrain of rats (Hoover & Vertes, 2007). The DBh projects heavily to the olfactory bulb, which has previously been identified as a potential source of ketamine-induced high-frequency oscillations (Hunt et al., 2019). Certainly, the biophysical basis of psilocybin-induced HFO remains enigmatic. 5-HT_2A_-R agonists and antagonists can induce increases and decreases in HFO power respectively, prompting a suggestion that HFOs may be modulated by 5-HT_2A_Rs (Goda et al., 2013). However, pyramidal and interneuronal expression levels of 5-HT_2A_Rs are lower in infralimbic cortex than either prelimbic or cingulate cortices (Santana & Artigas, 2017), while all three subregions express similar levels of 5-HT_1A_Rs (with highest expression in layer VI). Preferential effects of psilocybin on the infralimbic cortex may therefore reflect proportionally greater activation of 5-HT_1A_Rs at low doses. Indeed, the mPFC region investigated in this study corresponds to the region of highest 5-HT_1A_R expression across the whole cortex in rats, macaque monkeys and humans (Froudist-Walsh et al., 2023). As this is not true for 5-HT_2A_Rs (Froudist-Walsh et al., 2023; Palomero-Gallagher & Zilles, 2023), psilocybin‘s effects on mPFC could be more strongly driven by 5-HT_1A_R effects than other parts of cortex. These gradients of serotonin receptor densities across the cortex, with relatively higher 5-HT_1A_R expression in mPFC, hints at a potential mechanism through which top-down control may be modulated by psilocybin.

### Acute effects of psilocybin on cellular activity

Until now, very few studies quantified the effects of psilocybin on neural spiking *in-vivo*. Golden & Chadderton previously identified acutely increased population firing rates within the mouse anterior cingulate cortex after 2mg/kg psilocybin (Golden & Chadderton, 2022). Our contrasting finding of net decreases in population firing rates across the mPFC may reflect differences in species, dosage, behavioural state and/or sampling bias (Neuropixels enable wider sampling of pyramidal cell and interneuronal spike trains). Other studies have assessed the effects of DOI, another 5HT_2_ agonist, with mixed findings of either increased (Puig et al., 2003; Scruggs et al., 2003) or decreased (Ashby et al., 1989; Brys et al., 2023; Wood et al., 2012) neuronal activity within the rodent mPFC. Wood et al suggest these inconsistencies could be explained by the brain area and layer recorded from – studies that showed an increase after DOI were primarily from recordings of prelimbic and infralimbic layer V. In contrast, and similarly to our results, studies showing a decrease in firing after DOI were more heterogeneous in brain area and layers recorded (Wood et al., 2012). In addition, since DOI is not a significant agonist of 5HT_1A_Rs, our results also likely reflect the action of psilocybin on both 5HT_1A_ and 5HT_2A_ receptors.

There are several potential mechanisms for psilocybin-induced decreases in the spiking activity of pyramidal neurons. The subcellular distribution of 5-HT receptors suggests that the predominance of inhibitory 5-HT_1A_Rs on somatic, peri-somatic and axon initiation segments (AIS) might supress neural spiking, while high density of 5-HT_2A_Rs on distal dendrites can lead to stronger synaptic excitations and dendritic signal reception (Puig & Gulledge, 2011). Increased 5-HT_2A_R-dependent activation of potassium M-currents in mouse mPFC has also been reported to decrease the neuronal spiking through increased spike frequency adaptation and depolarization block (Ekins et al., 2023).

Although the net decreases in firing rate identified in our study belie a heterogenous mix of increased, decreased or unaltered firing rats in individual cells, they suggest that psilocybin leads to increased inhibition (and/or decreased excitation) within the mPFC, potentially suppressing the PFC’s top-down contributions of cognition during the drug’s acute influence. Whilst net suppression of PFC output is consistent with the broadband decrease in LFP power, we also identified an increase in the correlated activity between pairs of cells. Since at both doses of psilocybin, this increase was localised to pairs of cells within a region and not between regions, this finding supports psilocybin leading to a desynchronised state that favours local connectivity which may be reinforced by emergence of high-frequency oscillations (Brys et al., 2023; Smausz et al., 2022).

### Psilocybin-induced slowing and constraining of mPFC dynamics and influence

Several studies using indirect, non-invasive neuroimaging techniques (fMRI/MEG/EEG) have reported increased LZ complexity of brain dynamics as an empirical characteristic of altered consciousness in humans under a variety of psychedelic drugs, including psilocybin, LSD and DMT (Carhart-Harris et al., 2014; Lebedev et al., 2016; Schartner et al., 2017; Timmermann et al., 2019; Varley et al., 2020; Viol et al., 2017). Psychedelic-enhanced LZ complexity is also associated with more frequent recurrent state transitions in fMRI signals (Singleton et al., 2022). These studies mainly dealt with cortex-wide dynamics and low spatiotemporal resolution metrics of neural activity, computing LZ complexity by compression of multi-channel timeseries data from multiple cortical regions. Single channel analyses for individual cortical regions in MEG recordings under psilocybin have also shown increased LZ complexity, but the increase was focused within parietal-occipital regions, while the PFC was largely unaffected (Schartner et al., 2017).

Translating between fMRI and MEG/EEG signals primarily originating from slow time-scale membrane potential dynamics (such as synaptic events) and our analysis of psilocybin’s effects on mPFC spiking dynamics is complex. Nevertheless, our finding of decreased network complexity acutely following psilocybin was unexpected. The ‘relaxed beliefs under psychedelics’ (REBUS) hypothesis states that top-down signal transmission along the cortical hierarchy of signal processing is reduced while the bottom-up information transfer significantly increases (Carhart-Harris & Friston, 2019). This has been supported by neuroimaging studies examining the directed functional connectivity under psychedelic drugs (Girn et al., 2022; Shinozuka, Tewarie, et al., 2024). This may lead to increased synaptic activity in the higher associative cortical areas, causing increase in the fMRI and MEG/EEG signal diversity (Schartner et al., 2017). However, decreased mPFC neural spiking may also support the REBUS hypothesis, as the top-down input to lower cortical areas would be significantly reduced. This suggests that the locally observed decrease in mPFC network state transitions and LZ complexity play a significant role in flattening of the global cortical hierarchy.

### Delayed emergence of elevated gamma rhythms

Several studies have supported the notion that psychedelics like psilocybin may act as “psychoplastogens”, with their acute effects initiating circuit plasticity and enduring brain and behavioural changes (Heifets & Olson, 2024). The most prominent finding from our longitudinal data was a gradual increase in beta and slow gamma-frequency activity, that was highest on day 6 post-injection of 0.3mg/kg psilocybin. Gamma oscillations are generated by rhythmic activity in interconnected inhibitory-inhibitory or excitatory-inhibitory loops (Buzśaki & Wang, 2012), particularly involving fast-spiking parvalbumin (PV)-positive interneurons and their rhythmic entrainment of pyramidal neurons with feedback inhibition (Smausz et al., 2022; Sohal et al., 2009; Wang, 2010). The gradual increase in gamma shown in our study may therefore reflect psilocybin-induced plasticity in these microcircuits, ultimately leading to an alteration to E/I balance. Within the limited existing data on long-term effects of psilocybin, a change in gamma activity has not been investigated. In a study of resting state functional connectivity in 10 psychedelic naïve volunteers, McCulloch et al identified decreased connectivity within the prefrontal cortex 1 week (but not 3 months) after a ∼0.3mg/kg dose of psilocybin (McCulloch et al., 2022). Similarly, we found long-term decreases in broadband coherence across the CG/PL/IL cortices. This, combined with the finding of increased gamma power, corroborate previous findings of psychedelics inducing cortical desynchronisation (Muthukumaraswamy et al., 2013; Smausz et al., 2022).

### Limitations and conclusions

One challenge with analysing this data arose from changes in mPFC neurophysiology following saline vehicle injections, most likely reflecting effects of stress from the injection procedure. Since the injection procedure was consistent across drug and vehicle conditions and the order was randomised across animals, our analyses control for these effects. Nevertheless, lower-stress injection methods without restraint of the animal would be useful for future studies (Dearnley et al., 2021; Stuart & Robinson, 2015).

We have identified that a systemic injection of psilocybin leads to localised, state-dependent changes to the medial prefrontal cortex of rats. Our findings of decreased spectral power at low frequencies, increased power above 80Hz, decreased cell firing rates and signal complexity, local slowing of neural dynamics, and increased intra-regional cell pair correlations add subregional and cellular resolution to previous findings of psilocybin inducing desynchronisation across the brain. Several of these effects were most prominent during disengaged rest periods, consistent with clinical trial practice in which psilocybin is administered in low-stimuli environments. In addition, the localised infralimbic HFO could represent a novel biomarker of psychedelic action. Optogenetic modulation of infralimbic activity can recapitulate the enduring effects of ketamine on structural plasticity and behaviour (Fuchikami et al., 2015) and, in humans, Brodmann area 25 (the closest homolog to infralimbic cortex in rats (Nett & LaLumiere, 2021) has been associated with hyperactivity in treatment-resistant depression. A decrease in activity in this area has also been associated with clinical response to both antidepressant medication and deep brain stimulation (Mayberg et al., 2005). Our results complement these findings provoking further investigation into the infralimbic cortex and its circuitry as therapeutic targets preferentially modulated by psilocybin or more selective derivatives.

## ACKNOWLEDGEMENTS

This work was supported by a Wellcome Senior Research Fellowship in Basic Biomedical Science (202810/Z/16/Z), a BBSRC ‘Cognitive computational neuroscience’ award (BB/X013243/1) and by Compass Pathways. Our thanks to the University of Bristol Animal Services Unit staff for all their care and support.

## AUTHOR CONTRIBUTIONS

Conceptualization, MWJ and CWT; Methodology MWJ, CWT, CTG, RJP, SF-W and RG; Investigation RJP; Formal Analysis RJP, RG and SF-W; Data Curation RJP and RG; Writing – Original Draft RJP; Writing – Review & Editing RJP, RG, MWJ, SF-W, CWT and CTG; Visualization RJP and RG; Supervision MWJ and SF-W; Funding Acquisition MWJ and SF-W

## MATERIALS AND METHODS

### Animals and surgery

All procedures were performed in accordance with the UK Animals Scientific Procedures Act (1986) and were approved by the University of Bristol Animal Welfare and Ethical Review Board. Rats were housed individually in transparent, high-roofed cages (37x31x40cm lwh) to avoid damage to cranial implants; cages were enriched with wooden blocks and flowerpots. Water was provided *ad libitum* throughout, with food controlled to maintain body weight at no less than 85% of free-feeding values after recovery from surgery. Rooms were temperature- and humidity-controlled, with lights on at 08:15 and off at 20:15.

2 weeks after delivery to the unit, six adult male Lister Hooded rats (340-400g, 12-16 weeks of age, Charles River UK) underwent stereotaxic surgery under isoflurane recovery anaesthesia (4% induction, 1-2% maintenance), for implantation of Neuropixels 1.0 probes (IMEC, Belgium) into the medial prefrontal cortex (mPFC). Rats were each implanted with a Neuropixels probe into the mPFC (AP: +3.25-3.6mm; ML: 0.7-0.72mm; DV: -5.5mm) spanning the infralimbic cortex (IL), prelimbic cortex (PL) and cingulate cortex (CG). An i.p. injection of the anti-inflammatory dexamethasone (1mg/kg) and a local s.c. injection of the anesthetic Xylocaine 2% with adrenaline into the scalp was given prior to incision. Stainless steel screws (1.1mm diameter) were placed in the skull contralateral to the probe target location and across the skull for support. A further screw was placed overlying the cerebellum for a ground/reference site and Metabond (Sun Medical, Japan) was applied over all screws. Next, a ∼1.5 x 1.5mm craniotomy window was drilled into the skull for probe implantation. For a subset of animals (n=4) an automatic micromanipulator (IVM Mini Single, Scientifica, UK) was used to lower the probe into the brain at a speed of 0.2μm/s with manual lowering at a similar speed in the remaining animals. Once implanted, Vaseline was applied to cover the craniotomy and the remaining exposed shank of the probe. Dental acrylic with gentamycin antibiotic (DePuy Synthes, USA) was applied in a first layer over the surface of the skull, with further dental acrylic (Simplex rapid, Kemdent, UK) used for the remainder of the implant construction. A copper mesh cone encircling the implant was cemented around the surface of the skull with a ∼3cm end of a falcon tube cemented on top to contain and protect the electrodes and headstage. The cerebellar skull screw was connected to the shorted reference/ground on the Neuropixels probe and to the copper mesh cone via silver wires. A Neuropixels headstage was then connected to the probe and cemented to the side of the falcon tube before the lid was screwed on to protect the implant. An i.p. injection of the analgesic buprenorphine (0.05 mg/kg) was given post-surgery.

### Task protocol

An average of 13±4 days after surgery, rats were trained on an operant task. Training lasted for 6-7 days, outlined in Figure 1A. After rats were trained, they received an injection of either COMP360 (Compass Pathways’ investigational formulation of synthesized crystalline polymorph psilocybin) or vehicle (saline) counterbalanced across animals, with a 14-day washout period prior to receiving the alternative injection. Psilocybin was dissolved in saline at a concentration of 0.3 mg/ml. The drug and vehicle were intraperitoneally (i.p.) administered at a volume of 1 ml/kg in counterbalanced order across rats. The 0.3mg/kg dose was the focus of this study. It was chosen because it approximates the therapeutic dose used in clinical trials (Goodwin et al., 2022) and has been shown to be effective at long-term modulation of affective memory in a rat translational model (Hinchcliffe et al., 2024). However, in a subset of 4 rats a 1mg/ml solution was additionally administered 1ml/kg after a further 14-day washout period.

On the day of injection, recordings were first taken during a baseline rest and task block before a further 3 rest blocks and 2 task blocks were recorded after injection of the drug. Subsequent recordings were also taken during 2 rest and 2 task blocks on post-injection days 1, 2 and 6. Further details are outlined in Figure 1B. In all cases two rats were run through the protocol within the same day (one morning, one afternoon) with recordings starting at approximately the same time for each rat across days. All recordings were performed between 08:00-18:00, with drug injections between 09:00-10:00 for rats recorded in the morning, and 14:00-15:00 for rats recorded in the afternoon. Since two rats did not have baseline recordings prior to saline injections, final sample sizes for each condition were Saline: N=4, 0.3mg/kg: N=6, 1mg/kg: N=4.

Throughout the injection day and subsequent recording days, single unit activity and local field potentials (LFP) were recorded simultaneously using SpikeGLX software (Karsh *et al*., Janelia Research Campus). LFPs were sampled at 2.5 kHz; extracellular action potentials were sampled at 30 kHz with a 300Hz high pass filter. Video monitoring was collected simultaneously throughout the task and home-cage recordings.

At the end of experimental procedures, rats were deeply anaesthetised with sodium pentobarbital and transcardially perfused with 4% paraformaldehyde. 40µm brain sections were collected and immuno-stained with DAPI, GFAP, and IBA1 for staining cell nuclei, astrocytes, and microglia to identify glial scaring around the probe location. Histological images were mapped to a rat brain atlas (Paxinos & Watson, 2007) using the open-source python software HERBS (Fuglstad et al., 2023). Identification of the Neuropixels probe tip was then used to interpolate the location of the probe within the prefrontal cortex (Suppl. Fig.1).

### Analysis of local field potentials and single units

Data were processed in Matlab (Mathworks, MA) and Python (Python Software Foundation). Spectral power density from LFPs was calculated using the MATLAB pwelch function with 4s windows, no overlap, 0.25Hz frequency bins. A single LFP channel from each brain region was selected for further analyses (except for Figure 3, where all LFP channels were assessed). The infralimbic LFP channel was selected as the channel with the highest peak in 100Hz power within the infralimbic cortex (364±317µm [mean ± std] from the ventral boundary). Since recorded channels within the prelimbic cortex covered its entire dorsal-ventral boundary for all animals, the prelimbic LFP channel was selected from the approximate centre of the region (808±85µm from the ventral boundary). Since recorded channels within the cingulate cortex did not cover the entire dorsal-ventral boundary for all animals, the cingulate LFP channel was selected as the highest channel above the ventral boundary which was in an equivalent location across all animals (284±38µm from the ventral boundary).

For the spike data, a common average reference was applied across all channels. Single units were then isolated using automated clustering software (Kilosort 3; Pachitariu *et al*., 2021) followed by post-processing using the EcEphys pipeline (Allen Institute for Brain Science) and verification using Phy2 (Rossant *et al*., 2021). For post-processing, only units with isi violations <0.5, amplitude cutoff <0.1 and presence ratio >0.9 were selected for further analyses, following default criteria from AllenSDK. Putative pyramidal cells, narrow interneurons and wide interneurons were classified using CellExplorer (Petersen et al., 2021) based on their waveform spike width (<425μs for narrow interneurons) and tau rise in the autocorrelation (>6ms for wide interneurons). Cross-correlations were calculated with 2ms bin size and normalized by conversion to z-score. To avoid spurious correlations based on low activity units, only units with firing rates >0.15Hz during each 15minute block were included in the cross-correlation analyses, as similarly used in previous analyses (Kersanté et al., 2023).

### Mean-squared displacement and Lempel-ziv complexity of mPFC dynamics

For each 15-minute block, the spike trains of individual units were time-binned using a bin size of Δ*t* = 20 *ms*. For binarization, a time-bin was assigned 1 if it had one or more spike counts, and zero otherwise. Thus, the instantaneous state, ***S_t_***, of a neural network representing the mPFC region at time *t* was defined by a N-bit binary state, 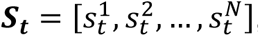, where 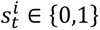 and *N* is the number of units recorded within the region. The mean-squared displacement, MSD, of the network state (Munn *et al.,* 2021) across a time step, *n*Δ*t*, ahead is given by,

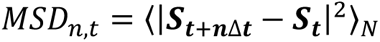

For each time *t*, we computed MSD values for multiple time steps of size *n* = 1 − 50. We then averaged the MSD values across n, *MSD*_*t*_ = 〈*MSD*_*n,t*_〉_*n*_ (Munn et al., 2021). This is equivalent to averaging the MSDs over 1 *sec* duration forward from each state. Given Δ*t* = 20*ms*, each 15min block contained temporally ordered 45,000 network binary states. Averaging the MSDs of jumps from each state to the next 50 states ahead meant discarding the final 50 states out of the 45,000. Thus, the mPFC network dynamics in an individual rat during each trial block a particular drug was characterized by a distribution over 44,950 MSD values.

Note that there were two post-injection blocks for each of the rest and task conditions. To examine how psilocybin impacts network MSD during rest and task blocks, we pairwise averaged the MSD values across the two post-injection blocks into one common post-drug MSD distribution. This was performed separately for rest and task blocks. Next, to account for the variability in baseline MSD distributions across rats, we z-scored the pre-drug distribution for each rat to have zero mean and unit variance. We also transformed the post-drug distribution for each rat, by subtracting the mean of the corresponding original pre-drug distribution from the post-drug distribution, which is further scaled by the standard deviation of the original pre-drug distribution. We then pooled the respective MSD values across rats into a single MSD distribution. Altogether, we had six post-drug pooled MSD distributions for the three drugs (saline, 0.3 mg/kg psilocybin, 1mg/kg psilocybin) and the two states (rest and task).

To compute the Lempel-Ziv (LZ) complexity of the mPFC network dynamics, we binarized the single unit spiking activity data using a time-bin of 10 *ms*. Using the original LZ complexity algorithm on finite sequences (Szczepański et al., 2004; Zhang et al., 2001), we computed the LZ complexity of binarized 15 minutes-long activity sequences of the individual units. Thus, for each drug and state, we obtained a unit-wise ordered list of LZ complexity values for the entire mPFC, by pooling the values across all cell types, mPFC sub-regions, and rats. Further, we pairwise averaged the LZ complexity values of the individual units across the two post-drug resting (task) states to obtain a single list of post-drug resting (task) LZ complexity values for each drug.

### Statistical analyses

All graphs plot means ± S.E.M unless otherwise stated. Boxplots show mean, median, 25^th^ and 75^th^ percentiles. LFP data is shown as an average across animals, single unit data is shown as an average across units. Changes in comparison to baseline were obtained by dividing the average post-injection values by the average baseline values for both drug and saline, conducted separately for rest and task conditions. The change after psilocybin was then divided by the change after saline to calculate percent differences between conditions. Depending on the data type, distributions, and variance, comparisons between saline and psilocybin were assessed using either parametric or non-parametric tests, including all three drug conditions (saline, 0.3mg/kg, and 1mg/kg psilocybin) where possible. Differences in behaviour, power density, and coherence were assessed with either a one-way anova (ANOVA1), two-way anova (ANOVA2), a repeated measures anova (RM-ANOVA), or a generalised linear model (GLM). Differences in single unit data were assessed with either a Scheirer-Ray-Hare test (SRH), a Kruskal-Wallis test (KW), a permutation test (PT), or a Chi-squared test (chisq). In all cases, if the test was significant, this was followed by a post-hoc pairwise comparison test (paired and unpaired T-test or Wilcoxon signed rank and rank sum test) followed by a Bonferroni correction (BC) for multiple comparisons. Since analyses of LFP data including power density and coherence analyses involves testing a substantial number of frequency variables (from 0 to 200Hz at 0.25Hz resolution), BC was not applied. Instead, a partial Bonferroni correction was used. For this, a principal component analysis was applied to the variable of interest and the number of output principal components that explained over 95% of the variance in the data was used as an the alpha correction factor, as previously described (i.e. α=0.05/number of principle components explaining >95% variance; (Scholtens et al., 2014)). Since the MSD and LZ complexity analyses were comprised of a large number of replicates with widespread significance, Cohens d was additionally calculated to gain insight into the strength of the statistical significance. Data from four rats were used in all analyses with data from the additional two rats used wherever possible (N=4 for all figures, unless otherwise stated).

