## Supplemental Figures 1-8 and Table 1 for "Short- and long-term reconfiguration of rat prefrontal cortical networks following single doses of psilocybin"

Ross J. Purple et al.

### Supplementary Figures

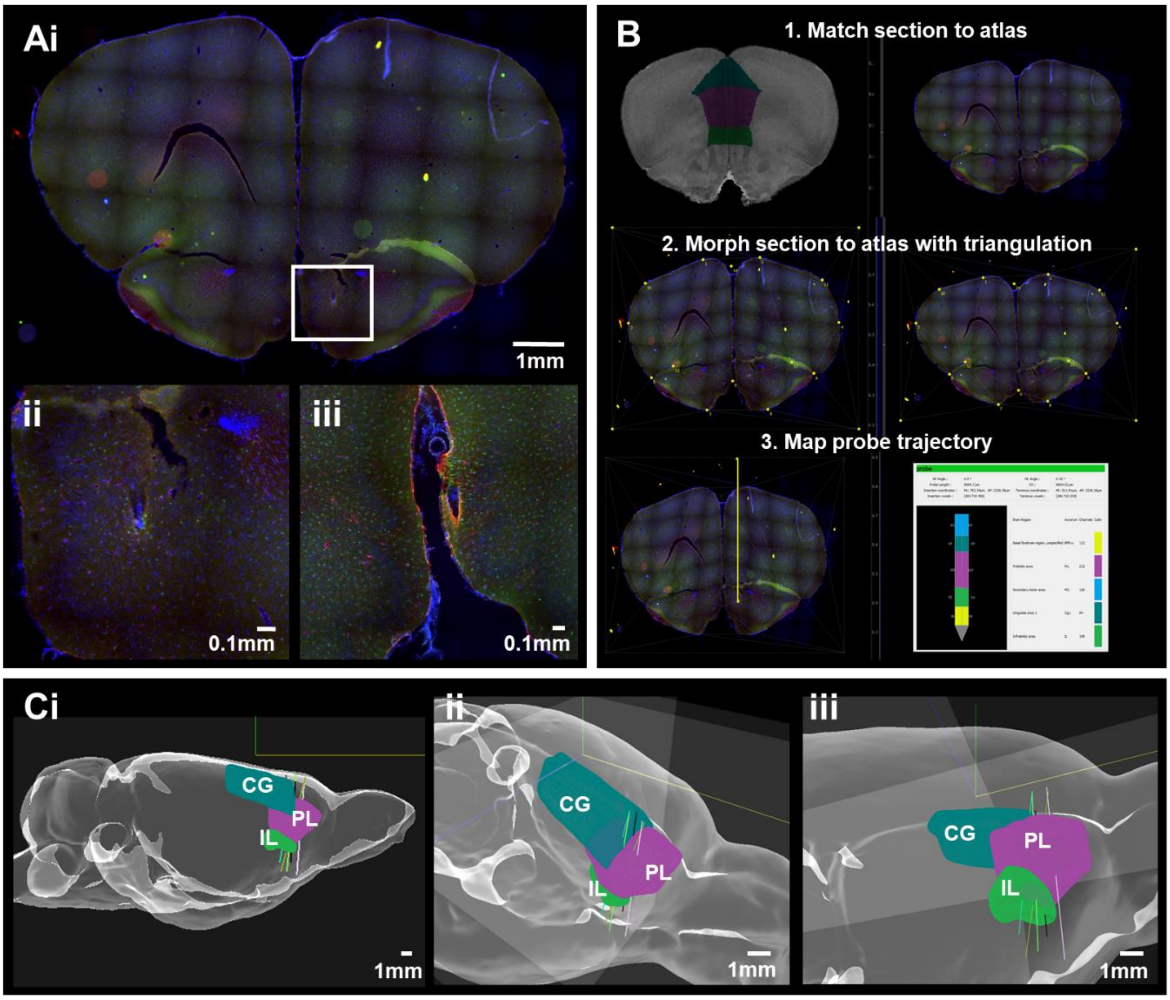


***Supplementary Figure 1:*** *Anatomical identification of the Neuropixels probe.* ***(A)*** *(i) 50um coronal brain slices were stained for glial markers (GFAP, IBA1) to identify scarring from Neuropixels probe. Sections containing scarring from the probe tip (centre of white square) were identified. (ii) Magnification of white square from section in panel A, showing scarring from the probe tip. (iii) A second example of probe tip from a separate rat.* ***(B)*** *Sections were processed using HERBS (Fuglstad et al., 2023) to map the probe trajectory within the brain.* ***(C)*** *(i) Sagittal, (ii) Dorsal-ventral, and (iii) ventral-dorsal views of all Neuropixels probes implanted within the medial prefrontal cortex (CG=cingulate cortex, PL=prelimbic cortex, IL=infralimbic cortex).*

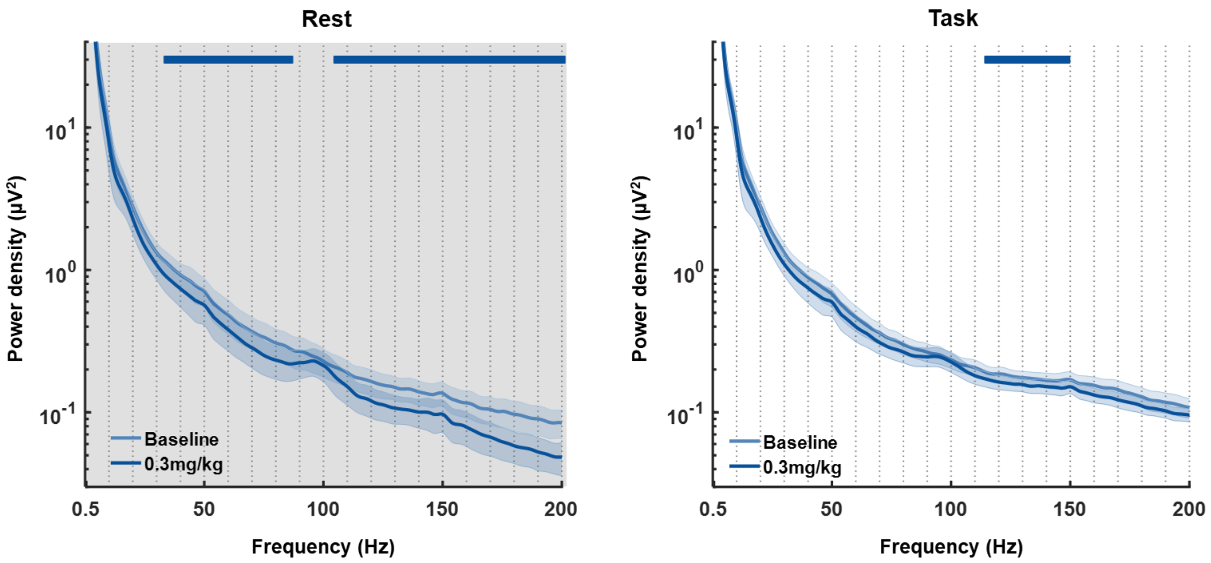


***Supplementary Figure 2.*** *Average power density from an infralimbic cortex LFP channel during a baseline and post-0.3mg/kg psilocybin injection rest block (left) and operant task block (right). Bars above trace represent significant differences between pre- and post-injection of 0.3mg/kg psilocybin (partial-Bonferroni corrected post-hoc t-tests, p<0.05). N=5.*

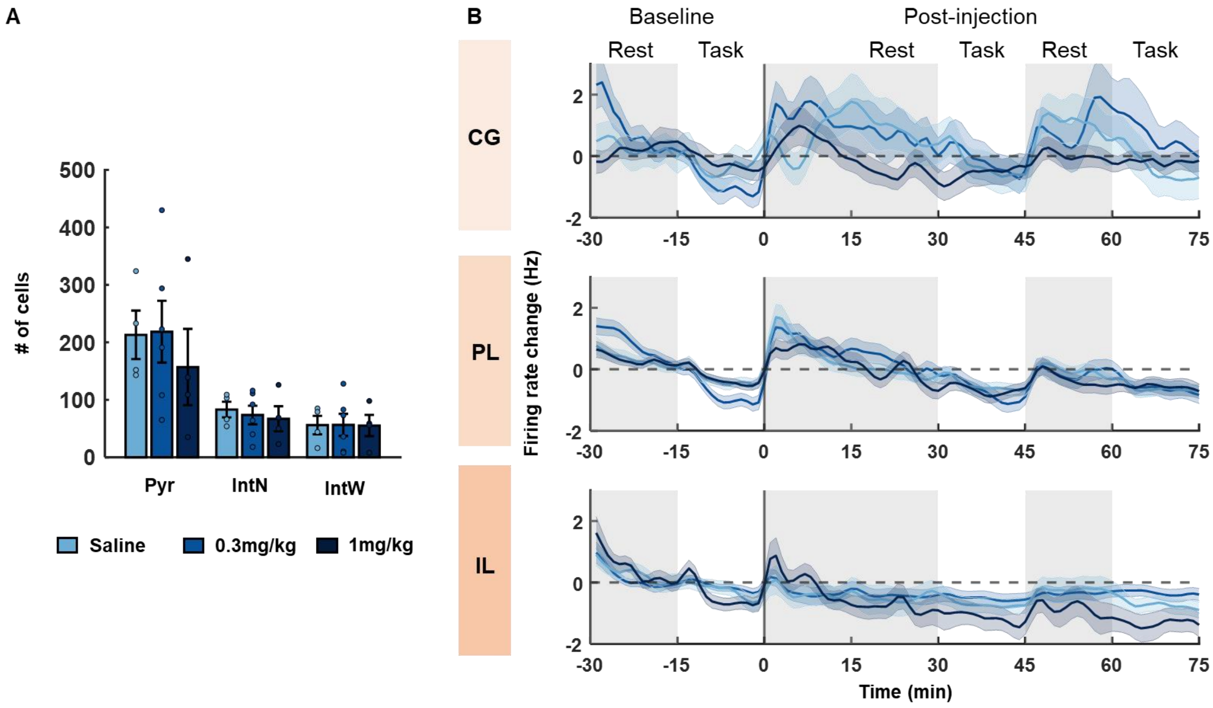


***Supplementary Figure 3: (A)*** *Total number of single cells identified during recordings on the day of injection of psilocybin or saline, separated into pyramidal cells (Pyr), narrow-waveform interneurons (IntN) and wide-waveform interneurons (IntW). Changes in interneuron cell firing.* ***(B)*** *Time-course of change in firing rates of interneurons in the cingulate cortex (CG; top), prelimbic cortex (PL; middle) and infralimbic cortex (IL; bottom) from pre-injection baseline. Firing rates are shown as a difference to the average firing rates during the 30-minute pre-injection baseline. Grey shading reflects rest blocks, white reflects task blocks. No differences were identified between conditions.*

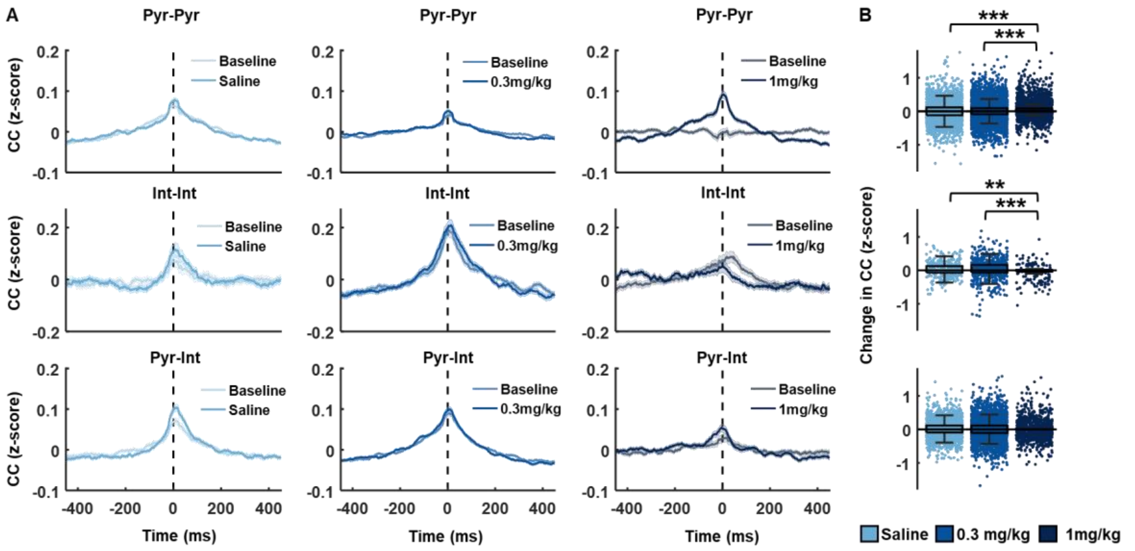


***Supplementary Figure 4:*** *Prelimbic cell cross-correlations.* ***(A)*** *Cross-correlated activity between cell pairs within the prelimbic cortex at baseline rest, and post-injection of saline (left, grey), 0.3mg/kg psilocybin (middle, light blue) and 1mg/kg psilocybin (right, dark blue). Shown are pyramidal-pyramidal (pyr-pyr), interneuron-interneuron (int-int) and pyramidal-interneuron (pyr-int) cell pair correlations.* ***(B)*** *Change in cross-correlated activity from baseline to post-injection rest blocks for pyr-pyr (top), int-int (middle) and pyr-int (bottom) cell pairs. A significant increase in cross-correlated activity was identified for pyr-pyr and int-int cell pairs after injection of 1mg/k psilocybin, compared to saline ($p<0.05$).*

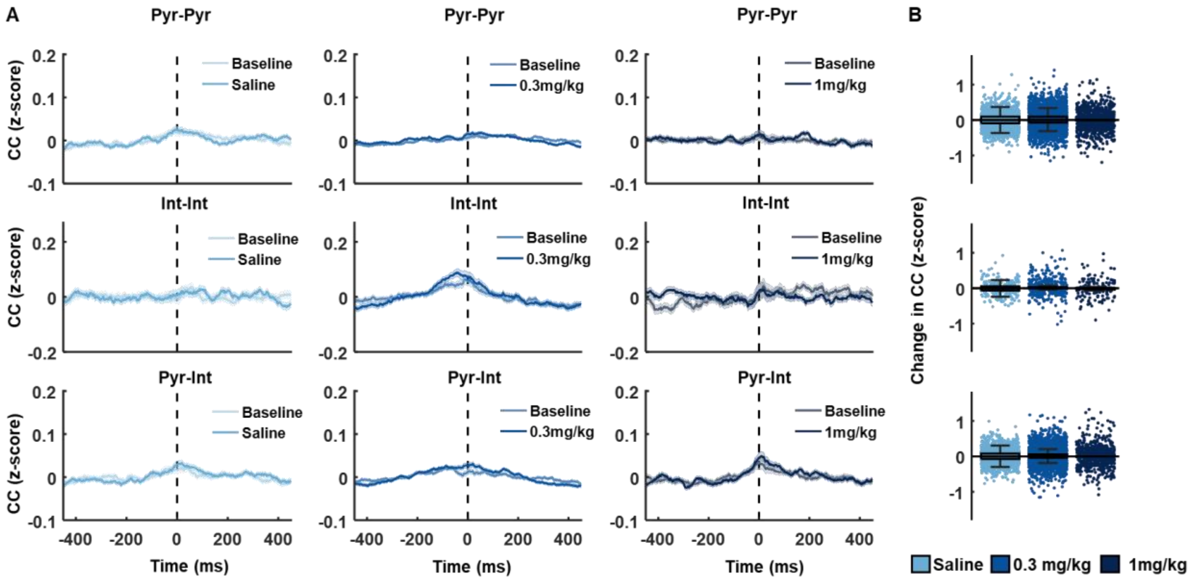


***Supplementary Figure 5:*** *Infralimbic-Prelimbic cell cross-correlations.* ***(A)*** *Cross-correlated activity between cell pairs across the infralimbic and prelimbic cortex at baseline rest, and post-injection of saline (left, grey), 0.3mg/kg psilocybin (middle, light blue) and 1mg/kg psilocybin (right, dark blue). Shown are pyramidal-pyramidal (pyr-pyr), interneuron-interneuron (int-int) and pyramidal-interneuron (pyr-int) cell pair correlations.* ***(B)*** *Change in cross-correlated activity from baseline to post-injection rest blocks for pyr-pyr (top), int-int (middle) and pyr-int (bottom) cell pairs. No differences were identified across conditions.*

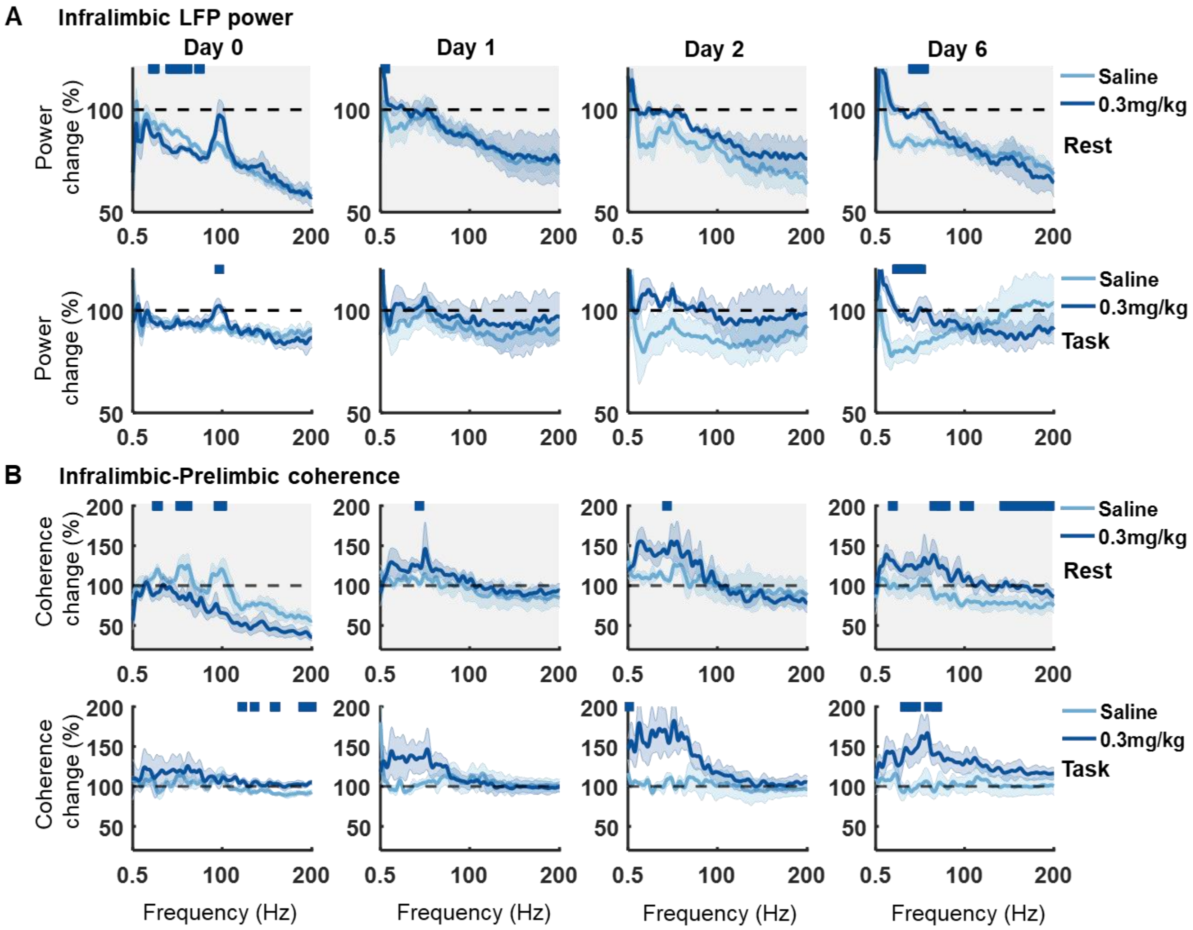


***Supplementary Figure 6: (A)*** *Change in power density from baseline to post injection of saline (light blue) and 0.3mg/kg psilocybin (dark blue) during rest (top, grey) and task (bottom, white) blocks, on the day of injection (Day 0) and subsequent days 1, 2 and 6 (all compared to day 0 baseline).* ***(B)*** *Change in infralimbic-prelimbic coherence from baseline to post injection of saline and 0.3mg/kg psilocybin across days during rest (top, grey) and task (bottom, white) blocks (all compared to day 0 baseline). Bars indicate significant differences between 0.3mg/kg psilocybin and saline (partial-Bonferroni corrected post-hoc t-tests, p<0.05). Shaded bands denote SEM.*

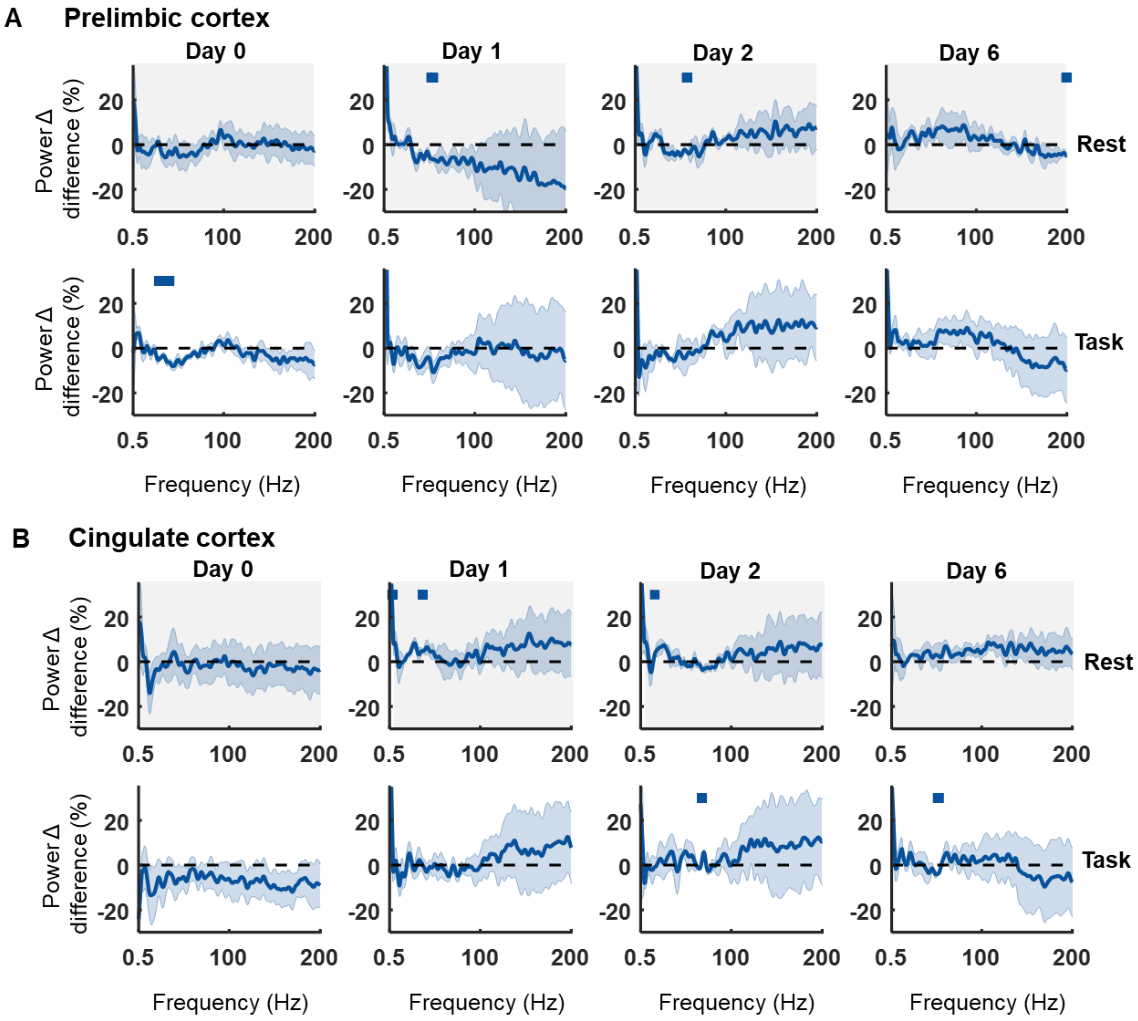


***Supplementary Figure 7:*** ***(A)*** *Difference between 0.3mg/kg psilocybin in comparison to saline in the change in power density within the prelimbic cortex from baseline to post injection during rest (top, grey) and task (bottom, white) blocks, on the day of injection (Day 0) and subsequent days 1, 2 and 6 (all compared to day 0 baseline).* ***(B)*** *Same but for LFPs in the cingulate cortex. Bars indicate significant differences between 0.3mg/kg psilocybin and saline (partial-Bonferroni corrected post-hoc t-tests, p<0.05). Shaded bands denote SEM.*

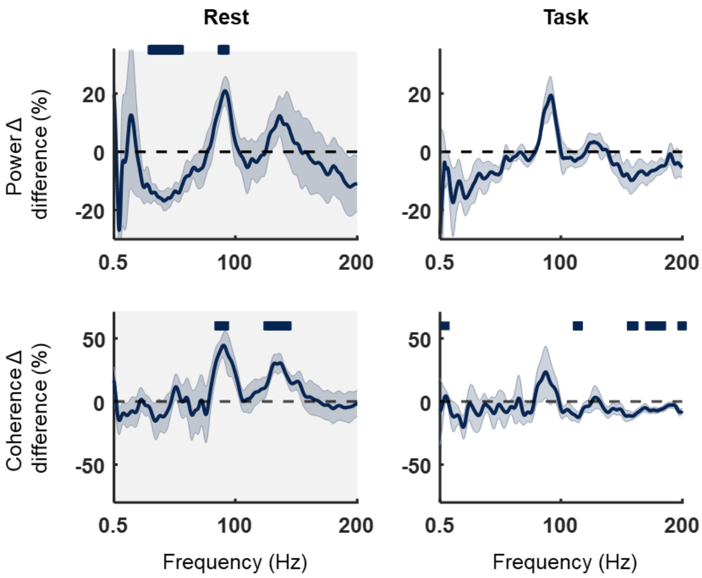


***Supplementary Figure 8:*** *Acute effects of 1mg/kg psilocybin on LFP power and coherence. (Top) Difference between 1mg/kg psilocybin in comparison to saline in the change in power density from baseline to post injection during rest (top, left) and task (top, right) blocks, on the day of injection (Day 0). (Bottom) Difference between 1mg/kg psilocybin in comparison to saline in the change in infralimbic-prelimbic coherence from baseline to post injection rest (bottom, left) and task (bottom, right) blocks. Bars indicate significant differences between 1mg/kg psilocybin and saline (partial Bonferroni corrected post-hoc t-tests, $p<0.05$).*

***Supplementary Table 1:*** *Two-way analysis of variance results for difference between saline and 0.3mg/kg psilocybin in the change in power and coherence from pre-injection baseline to post-injection day 1 and 2. N=4.*

| Block | Day | Factor 1 | | Factor 2 | | Interaction | |
|---|---|---|---|---|---|---|---|
| Power | | Frequency | | Drug | | Frequency x Drug | |
| Rest | 1 | F(800,4806)=3.37 | p<0.001 | F(1,4806)=67.98 | p<0.001 | F(800,4806)=0.36 | p=1.000 |
| Task | 1 | F(800,4806)=0.64 | p=1.000 | F(1,4806)=143.90 | p<0.001 | F(800,4806)=0.45 | p=1.000 |
| Rest | 2 | F(800,3204)=3.60 | p<0.001 | F(1,3204)=613.30 | p<0.001 | F(800,3204)=0.16 | p=1.000 |
| Task | 2 | F(800,3204)=0.52 | p=1.000 | F(1,3204)=566.35 | p<0.001 | F(800,3204)=0.14 | p=1.000 |
| Coherence | | Frequency | | Drug | | Frequency x Drug | |
| Rest | 1 | F(800,4806)=2.03 | p<0.001 | F(1,4806)=264.56 | p<0.001 | F(800,4806)=0.27 | p=1.000 |
| Task | 1 | F(800,4806)=0.95 | p=0.847 | F(1,4806)=159.96 | p<0.001 | F(800,4806)=1.06 | p=0.120 |
| Rest | 2 | F(800,3204)=2.69 | p<0.001 | F(1,3204)=113.85 | p<0.001 | F(800,3204)=0.76 | p=1.000 |
| Task | 2 | F(800,3204)=1.93 | p<0.001 | F(1,3204)=1352.7 | p<0.001 | F(800,3204)=1.41 | p<0.001 |
